# PerturbPlan: An analytical framework for designing Perturb-seq experiments

**DOI:** 10.64898/2026.05.22.727199

**Authors:** Ziang Niu, Yihui He, James Galante, Andreas R. Gschwind, Judhajeet Ray, Lars M. Steinmetz, Jesse M. Engreitz, Eugene Katsevich

## Abstract

CRISPR screens with single-cell RNA-seq readouts provide a powerful tool for characterizing the functions of noncoding elements and genes. However, designing these experiments to balance statistical power and cost is challenging, given the large number of design parameters. The only available tool for this purpose is a simulation-based power calculator, but it is computationally costly and requires high-performance computing to run. We derive a novel analytical formula for the power to detect perturbation-expression associations, recapitulating power estimates from the simulation-based tool while reducing runtime by up to seven orders of magnitude. This acceleration unlocks the possibility of interactive single-cell CRISPR screen design. Accordingly, we develop PerturbPlan, an interactive web application built on the analytical power formula. PerturbPlan helps users address 11 design questions for two types of single-cell CRISPR screens, Perturb-seq and targeted Perturb-seq (TAP-seq). We apply PerturbPlan to carry out a comparative analysis of three recent Perturb-seq designs, demonstrating how optimal design varies across experiments of different scales. We also use PerturbPlan to quantify the cost savings of a recent TAP-seq study relative to a hypothetical Perturb-seq study assaying the same perturbations, illustrating how the tool can inform decisions about targeted versus whole-transcriptome readouts. In sum, PerturbPlan is the first tool to facilitate flexible and interactive design of well-powered single-cell CRISPR screen experiments.

## Introduction

CRISPR screens with single-cell RNA-seq readouts^1–6^ (henceforth single-cell CRISPR screens) have become a staple tool for mapping regulatory relationships at genome scale. Versions of this assay include Perturb-seq^2^ and targeted Perturb-seq (TAP-seq),^6^ which read out the entire transcriptome or a chosen subset of genes, respectively. These assays have been applied with enhancer-targeting perturbations to map enhancers to their target genes,^6–9^ a critical step in tracing the path from variant to function for genetic loci associated with complex diseases.^10^ They have also been applied with gene-targeting perturbations to reconstruct gene regulatory networks.^2,11–12^ The resulting datasets are valuable not just for their biological insights, but also for training and benchmarking predictive models extending these insights to new perturbations and biological contexts. Such efforts have been undertaken in the context of both enhancer-targeting^13,14^ and gene-targeting perturbations,^15–18^ and are central to the recent push to use AI to build the virtual cell.^19,20^ Given the promise of single-cell CRISPR screens, these assays have played a central role in recent scientific consortia, including ENCODE4,^21^ the Impact of Genomic Variation on Function Consortium^22^ and Molecular Phenotypes of Null Alleles in Cells Consortium.^23^

For single-cell CRISPR screens to effectively facilitate biological discovery and predictive model training, it is critical that they achieve sufficient statistical power to detect gene expression changes in response to perturbations. Such tests are a basic unit of analysis conducted on single-cell CRISPR screen data, and their results often form the basis for downstream analyses, including training predictive models. A recent concern in enhancer-gene predictive modeling illustrates the importance of power: enhancer-gene pairs can be erroneously labeled as having no regulatory effect due to low power rather than true absence of such a relationship, hampering model training and evaluation.^13^ However, a recent study found that small effect sizes (5-10%) are common among enhancer perturbations, but previous Perturb-seq screens are underpowered to detect such effects.^9^ For example, the Perturb-seq screen of Gasperini et al.^7^ was estimated to be well-powered to detect only about 10% of element-gene pairs with such small effect sizes. Underpowered experiments often detect mainly strong effects or those involving highly expressed genes, illuminating only a fraction of the regulatory landscape. In recognition of such issues, the data underlying the 2025 Virtual Cell Challenge was purposefully designed for high coverage in terms of cells per perturbation and reads per cell, though without a power analysis to quantify what effects could reliably be detected.^20^

Despite the importance of experimental design for single-cell CRISPR screens, significant unresolved challenges hamper this task in practice. One challenge is the diversity of experimental and analysis choices across lentiviral delivery, library preparation, sequencing, and differential expression testing (**Fig. 1a**). Some choices are widely encountered in genomics, like how many cells to profile or how deeply to sequence each cell, while others are specific to single-cell CRISPR screens, like how many genomic targets (e.g., enhancers or genes) to perturb. These choices impact not just the power of a single-cell CRISPR screen experiment but also its cost, and experimentalists may struggle to juggle these decisions (**Fig. 1b**), particularly because no dedicated interface for single-cell CRISPR screen design is currently available. The core technical problem is mapping all the parameters of an experiment to its power. The only existing approach to this problem is simulation-based.^9^ However, the rapidly increasing scale of single-cell CRISPR experiments^11^ makes simulation-based approaches computationally expensive or even infeasible. Indeed, the large number of parameters impacting power leads to a high-dimensional design space, meaning that a large number of parameter combinations must often be explored to arrive at an optimal experimental design. While an analytical power analysis framework exists in the adjacent domain of single-cell differential expression,^24^ this tool does not accommodate aspects unique to single-cell CRISPR screens and supports limited design problems (see “Comparison of PerturbPlan to scPower” section in the **Supplementary Note**). In sum, there is currently a pressing need for tools to guide the trade-off between power and cost in designing single-cell CRISPR screens.

**Figure 1.**
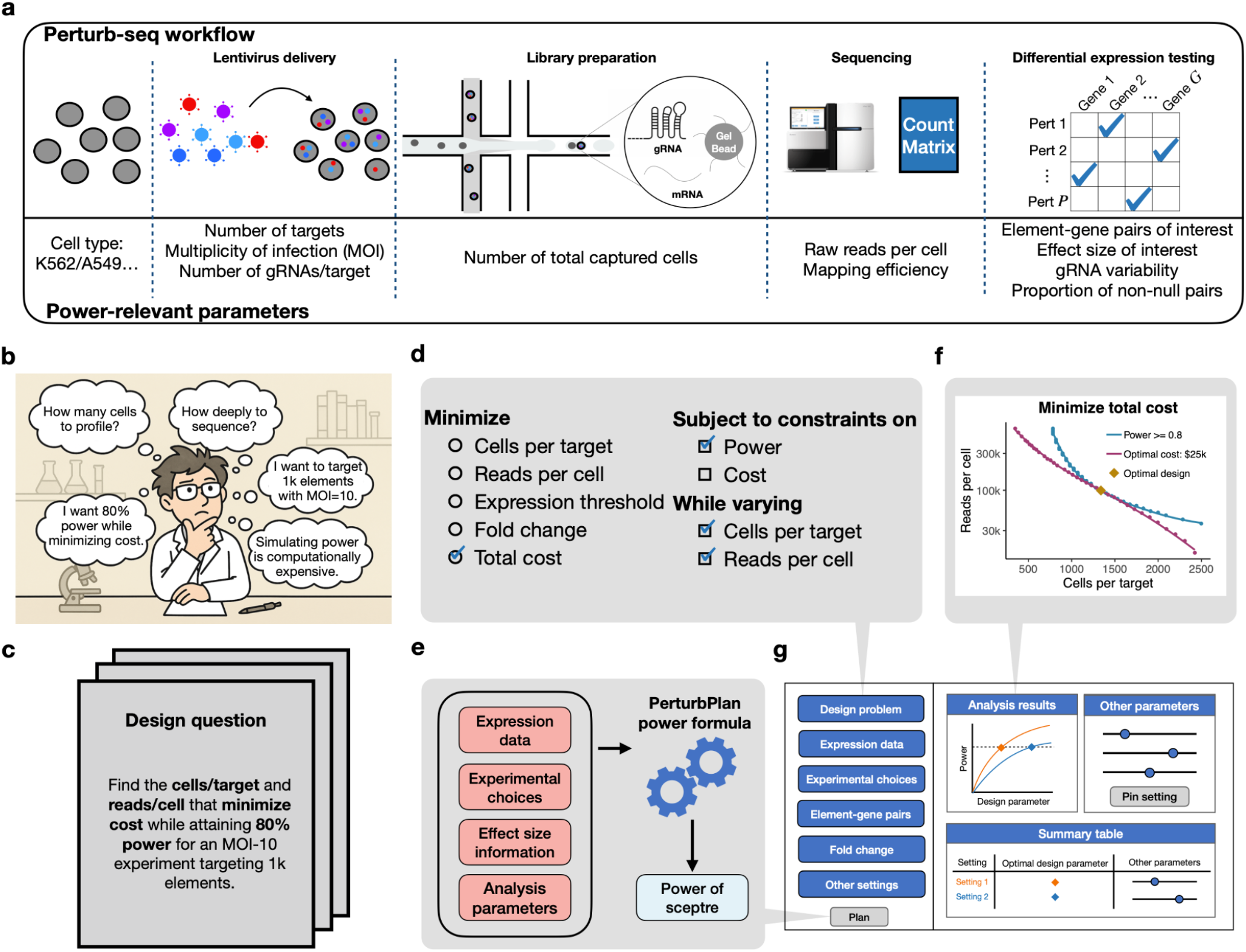
PerturbPlan provides a principled and flexible framework for single-cell CRISPR screen experimental design, supported by a user-friendly web application. **a:** Perturb-seq workflow and parameters relevant to power to detect perturbation-gene effects. **b:** Perturb-seq experimental design is challenging: it involves a number of design choices and constraints that are difficult to balance. **c:** Perturb-seq design problems can be formulated in a number of ways. The example given formalizes the design question suggested by panel **b**. **d:** A framework for specifying experimental design problems. The web app supports 11 design problems, each of which can be obtained by choosing one of five variables to minimize, whether to constrain just power or both power and cost, and whether to additionally vary cells per target, reads per cell, or both. The checked variables correspond to the design problem in panel **c**. **e:** PerturbPlan formula schematic. The salmon boxes denote user inputs and arrows indicate the direction of information flow. **f:** An example of a plot generated by PerturbPlan corresponding to the design setup in panels **a** and **b**. The points on the blue line have equal power 0.8. The points on the red line have equal cost $25k, which is the minimum cost across all the configurations on the blue line. The diamond indicates the optimal design. **g:** A schematic of the PerturbPlan web application, including sidebar for user input at left, and main panel for app output and user interaction at right.

To address this need, we present **PerturbPlan**, an analytical framework and interactive web application (www.perturbplan.com) for designing Perturb-seq and TAP-seq experiments. We resolve the core technical problem of power calculation by developing a novel analytical formula approximating the power of sceptre,^28,29^ a state-of-the-art tool for testing perturbation-expression association in single-cell CRISPR screens. This formula overcomes the computational limitations of simulation-based power calculation, enabling millisecond-speed power calculation for each parameter configuration and, in turn, real-time experimental design recommendations. We also develop the PerturbPlan web application, embedding the power formula within an experimental design framework addressing 11 common design problems. This application allows users to interactively explore design scenarios and tailor recommendations to their specific experimental context with the help of intuitive visualizations. We demonstrate that the PerturbPlan formula provides an excellent approximation to the output of the simulation-based tool while reducing its sequential runtime by seven orders or magnitude. In real-data downsampling benchmarks from Perturb-seq and TAP-seq datasets, PerturbPlan captured empirical power trends across design parameters, with empirical power meeting or exceeding predictions when the tool was applied based on a minimum effect size of interest. We apply PerturbPlan to (1) carry out a comparative analysis of three recent Perturb-seq designs, demonstrating how optimal design varies across experiments of different scales and (2) quantify the cost savings of a recent TAP-seq study compared to a hypothetical Perturb-seq study assaying the same perturbations, illustrating how the tool can inform decisions about targeted versus whole-transcriptome readouts. In sum, we provide the first interactive tool to systematically guide the design of single-cell CRISPR screen experiments.

## Results

### PerturbPlan supports flexible and interactive single-cell CRISPR screen experimental design

PerturbPlan is a power calculation engine wrapped in an intuitive interface for specifying design problems, uploading pilot data, configuring experimental choices, and visualizing analysis results. The user experience is driven by the design problem (**Fig. 1c**), a precise statement of the task at hand. We developed a structured framework for design problem specification (**Fig. 1d**) supporting 11 design problems (**Extended Data Tab. 1**) covering five choices of optimization targets (cells per target, reads per cell, expression threshold, fold change, and total cost), two choices of variables that can be constrained (power and/or cost), and two additional free variables (the numbers of cells per target and/or reads per cell). The tool can be used for any cell type and stimulation condition, provided that matching reference expression data are available. In particular, both the expression count matrix and the number of reads supporting each UMI are required (the latter is available, for example, from Cell Ranger’s standard output file `molecule_info.h5`). Reference expression data are built into PerturbPlan for several commonly used cell types; as the time of publication, these are K562,^7^ A549,^30^ iPSC,^31^ iPSC-derived neurons,^31^ CD8+ T cells,^32^ and THP-1 cells^33^ (**Extended Data Fig. 1**). PerturbPlan also supports custom reference datasets; functions needed for preprocessing raw expression data are available in the companion R package `perturbplan` (https://katsevich-lab.github.io/perturbplan/), including dedicated functions for Cell Ranger output. The power calculation engine (**Fig. 1e**) combines the reference expression data with experimental choices, analysis choices, and effect size information to obtain an approximation for the power of `sceptre`. The web application calculates the optimal solution to the design problem by searching over grids of parameters, computing power and/or cost for each. The result of this process is displayed as an intuitive visualization (**Fig. 1f**). The user interface (**Fig. 1g, Extended Data Fig. 2**) includes user input in a sidebar on the left and the visualization and recommended designs in the main panel on the right. Also included in the main panel are sliders allowing users to adjust key parameters; the speed of the power formula allows real-time updates for smooth interactivity.

PerturbPlan implements a statistically rigorous workflow for approximating the power of `sceptre` (**Fig. 2**). This workflow consists of five main steps. In the first step (**Step 1** in **Fig. 2**), we estimate baseline expression distributions on the reference expression data by fitting a negative binomial (NB) regression model to obtain relative expression and dispersion parameters for each gene. In the second step (**Step 2** in **Fig. 2**), we fit a sequencing saturation curve on the reference expression data, mapping reads per cell to library size (average UMIs per cell), using the preseqR package.^34,35^ In the third step, we derive an analytical distributional approximation for the test statistic of each hypothesis using the estimated baseline expression and library size from the first two steps (**Step 3** in **Fig. 2**). The test statistic analyzed is a simplified variant of the `sceptre` test statistic that adjusts for library size but not other covariates (**Supplementary Note**). We reasoned that, in homogeneous cell lines often used for single-cell CRISPR screens, library size is the most important per-cell covariate to account for. Furthermore, we expect that excluding batch effects during experimental design while including them during analysis would usually lead to conservative design. The fourth step provides a prospective analytical approximation of the Benjamini-Hochberg p-value threshold (**Step 4** in **Fig. 2**). This involves finding the root of a univariate function via an efficient bisection algorithm, using the distributional approximation from **Step 3**, the non-null proportion, and the target FDR level. Finally, in the fifth step (**Step 5** in **Fig. 2**), we compute overall power as the average power across individual element–gene tests, evaluated at the approximate p-value threshold from **Step 4**. The formula uses a normal tail probability rather than the resampling-based tail probability employed by `sceptre`. The difference between these two is more pronounced for lowly expressed genes, but we reasoned the difference would diminish upon averaging across genes spanning a range of expressions.

**Figure 2.**
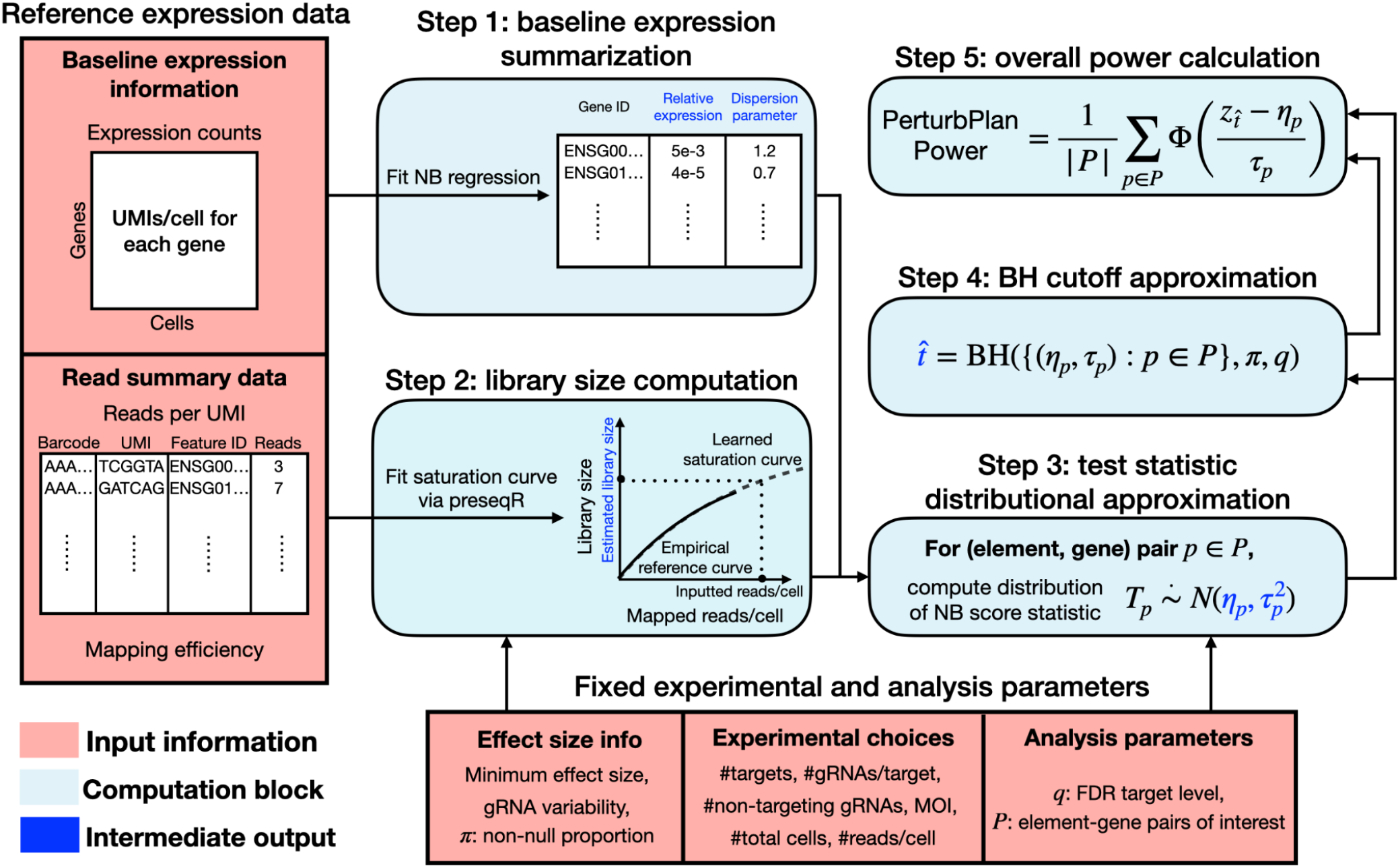
PerturbPlan power calculation workflow. **Reference expression data:** Baseline expression and read summary information for the cell type of interest. The reference expression data includes the gene expression count matrix, the number of reads supporting each UMI, and the mapping efficiency. **Fixed experimental and analysis parameters:** Information on effect sizes, experimental choices and analysis parameters. **Step 1:** A negative binomial (NB) regression is fit to each gene’s expression counts, yielding a per-gene baseline expression distribution parameterized by its relative expression and dispersion. **Step 2:** The reads per UMI data is used to fit a saturation curve via preseqR.^34,35^ The number of sequenced reads per cell is multiplied by the mapping efficiency to get the expected number of mapped reads per cell, which is plugged into the fitted saturation curve to give the expected library size, i.e., expected number of UMIs per cell. **Step 3:** A distributional approximation is obtained for negative binomial score statistics testing each element-gene pair using the input from **Step 1, Step 2** and **fixed experimental and analysis parameters**. The exact formula can be found in the **Methods** section. **Step 4:** An approximation of the BH p-value threshold is obtained from the distributional approximation, FDR target level as well as the non-null proportion parameter. The exact formula can be found in the **Methods** section. **Step 5:** Overall power is computed by averaging the detection power over all element-gene pairs of interest. We present an instance of the formula based on a left-sided test geared to detect perturbation-induced decreases in gene expression.

### PerturbPlan benchmarking and validation

We validated PerturbPlan using both synthetic and real data. We first assessed the speed and accuracy of its power estimates in a proof-of-concept simulation study (**Fig. 3a–b**), benchmarking against an existing simulation-based pipeline^9^ to estimate the power of `sceptre` for perturbation–gene testing in Perturb-seq. Across a range of configurations for cells per target (**Fig. 3a**) and fold changes (FCs) (**Fig. 3b**), the discrepancy between PerturbPlan’s estimates and the simulated power remains below 0.02. The analytical nature and software optimizations of PerturbPlan enabled sub-millisecond power computation per design configuration, whereas the simulation-based pipeline was approximately seven orders of magnitude slower in terms of CPU-hours, though wall clock runtime was somewhat faster due to parallelization (**Fig. 3c**).

**Figure 3.**
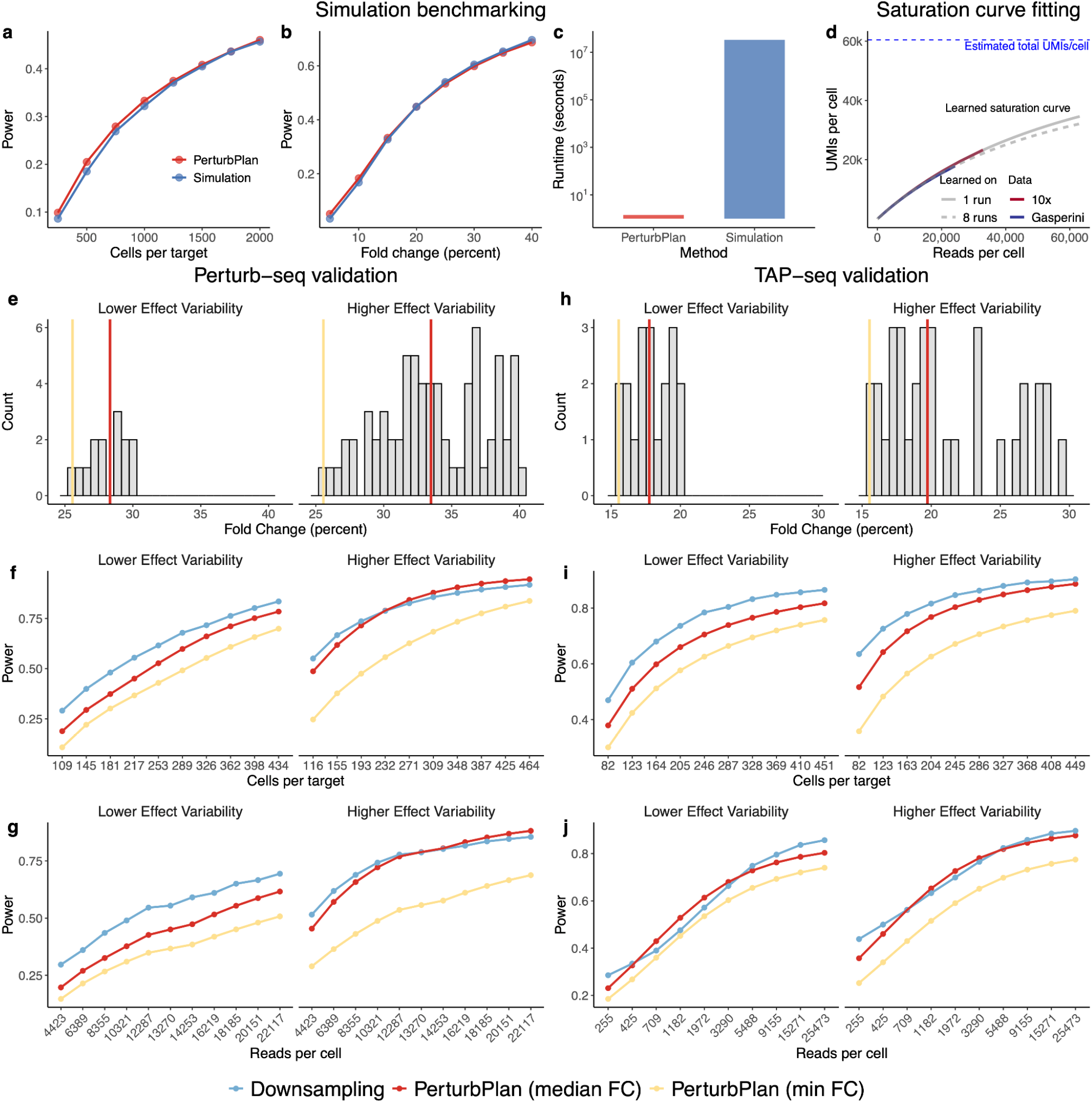
PerturbPlan recapitulates simulation-based power while being orders of magnitude faster, and provides reliable power estimates in real data validation studies. **a-c:** Simulation benchmarking with PerturbPlan. Power is compared across different values of cells per target and FC. In panel **c**, sequential CPU runtime to compute power for 1000 parameter configurations is compared between PerturbPlan and the simulation-based approach. **d:** Learning the sequencing saturation curve via preseqR. The dashed grey curve is the learned sequencing saturation curve based on 8 runs from Perturb-seq data,^7^ aligning closely with the solid curve learned only with 1 out of 8 runs. The curves both fit the downsampled curve used for learning (blue) and generalize well to independent Perturb-seq data^36^ on the same K562 cell type (red). **e-g:** Real data validation with Perturb-seq experiment^7^ for K562 cell type. We consider two different ranges of effect sizes (**e**) for positive controls. Power is compared across different configurations of cells per target (**f**) and reads per cell (**g**). We consider two estimates of effect size when applying PerturbPlan: one based on the median of the effect size distribution and one based on the minimum effect size. **h-j:** Real data validation with DC-TAP-seq^9^ for K562 cell type, similar to panels **e**-**g**.

We next evaluated PerturbPlan on real data. As a first step, we examined the performance of its preseqR-based saturation curve fitting model, which had previously not been applied to single-cell CRISPR screen data. As a basic check, we compared the fitted saturation curve to the empirical saturation curve from the Gasperini et al.^7^ (K562) and Shifrut et al.^32^ (CD8+ T) Perturb-seq datasets and found excellent agreement (**Fig. 3d, Extended Data Fig. 3a,b**). In particular, we found a better fit than that provided by the analogous component of scPower. For speed purposes, PerturbPlan uses only one sequencing run to fit the saturation curve. We verified that the saturation curve fit based on one run matched well with a saturation curve based on eight runs. Next, we assessed extrapolation performance from more shallowly sequenced reference datasets (**Extended Data Fig. 3c-f**) by downsampling reads in the Gasperini et al. and Shifrut et al. datasets. We found that the saturation curve fit remained stable even after approximately a 100-fold decrease in sequencing depth. Finally, we assessed extrapolation performance across datasets, leveraging a dataset produced by 10x Genomics in the same cell type, K562.^36^ The saturation curve fit to the Gasperini data matched well with the empirical saturation curve for the 10x Genomics data, including at sequencing depths beyond Gasperini’s (the 10x Genomics data was more deeply sequenced) (**Fig. 3d)**. This finding demonstrates that saturation curves based on preseqR can be successfully transferred across sequencing libraries based on different numbers of cells, which had not been previously investigated.^37^ In sum, we found that the PerturbPlan’s preseqR-based saturation curve fit demonstrated excellent fit and extrapolation.

We then validated PerturbPlan’s power estimates on enhancer-to-gene mapping screens using Perturb-seq^7^ and DC-TAP-seq.^9^ A key challenge in obtaining ground truth power based on real data is that, unlike in simulations, most experiments are conducted only once and are not replicated under identical parameter configurations. To address this, we developed a downsampling framework to mimic experimental replication (see **Methods**). Within this framework, we restricted attention to positive and negative control pairs so that the identities of the regulatory relationships among the pairs tested were known. We considered two ranges of effect sizes in each experiment (**Fig. 3e,h**) to cover weaker-effect and stronger-effect regimes. PerturbPlan requires as input the effect size and baseline relative expression. We obtain these by applying sceptre to the full dataset. Since the true effect sizes vary across pairs while PerturbPlan requires a single effect size as input, we applied PerturbPlan in two ways: using the minimum FC (for conservative power estimates) and the median FC (for more accurate power estimates). Our main findings from the real-data validation (**Fig. 3f–g, i–j**) are as follows. When using the minimum FC, PerturbPlan consistently yields power estimates that lower-bound the empirical power obtained from downsampling, confirming that the method provides conservative power estimates when supplied with the minimum effect size of interest. The gap between the estimated and empirical power grows as the effect size distribution becomes more dispersed (“Lower Effect Variability” versus “Higher Effect Variability”). When using the median FC, PerturbPlan’s power estimates more closely track the downsampled power but no longer guarantee a lower bound. We recommend PerturbPlan users specify the minimum FC of interest for conservative experimental design.

To assess the robustness of our power formula to the choice of reference expression data within the same biological system, we compared power estimates obtained from two Perturb-seq datasets generated in the K562 cell line. We verified that the broadly similar mean and dispersions of gene expression across the two datasets (**Extended Data Fig. 4a,b**) gave rise to consistent power estimates (**Extended Data Fig. 4c,d**) as well as similar optimal designs (**Extended Data Fig. 4e-h**).

### Power and cost comparison across three existing Perturb-seq designs

We applied PerturbPlan to compare the designs of three influential Perturb-seq studies—Morris^8^, Gasperini^7^, and Replogle^11^—and to compare the scales of optimally redesigned experiments for each. These studies were all carried out in K562 cells, but had different scientific goals and scales: Morris sought to characterize ~600 enhancers implicated by GWAS in blood traits, Gasperini to characterize ~6,000 active enhancers genome-wide, and Replogle to characterize all ~10,000 genes expressed in K562. Potentially due to cost constraints, the depth of profiling (in terms of reads per cell and cells per target) across studies decreased as the number of targets increased (**Fig. 4a**). This is particularly true for the Replogle study, in part because this was the only low multiplicity of infection (MOI) study among the three. Inputting the profiling depth and other parameters (**Methods**) into PerturbPlan, we computed the power of each study to detect a 25% FC (**Fig. 4b**). Since the inclusion of lowly expressed genes decreases power, we considered power among pairs subject to differing TPM thresholds. We also carried out a parallel analysis with no TPM threshold but varying the FC (**Fig. 4c**). Across both comparisons, we found that the Morris study had the highest power, the Gasperini study had somewhat lower power, and the Replogle study was substantially less powerful than the other two. For example, at a 25% fold change with all expressed genes (Figure 4c), the Morris, Gasperini, and Replogle studies achieved overall powers of 0.79, 0.62, and 0.24, respectively. These differences were driven primarily by the number of cells per target: approximately 1,000 for both Morris and Gasperini, but only approximately 200 for Replogle, which targets a much larger number of perturbations (10K vs. 600 and 6K). This ordering mirrors the orderings of reads per cell and cells per target (**Fig. 4a**), highlighting the central role of these quantities in determining power.

**Figure 4.**
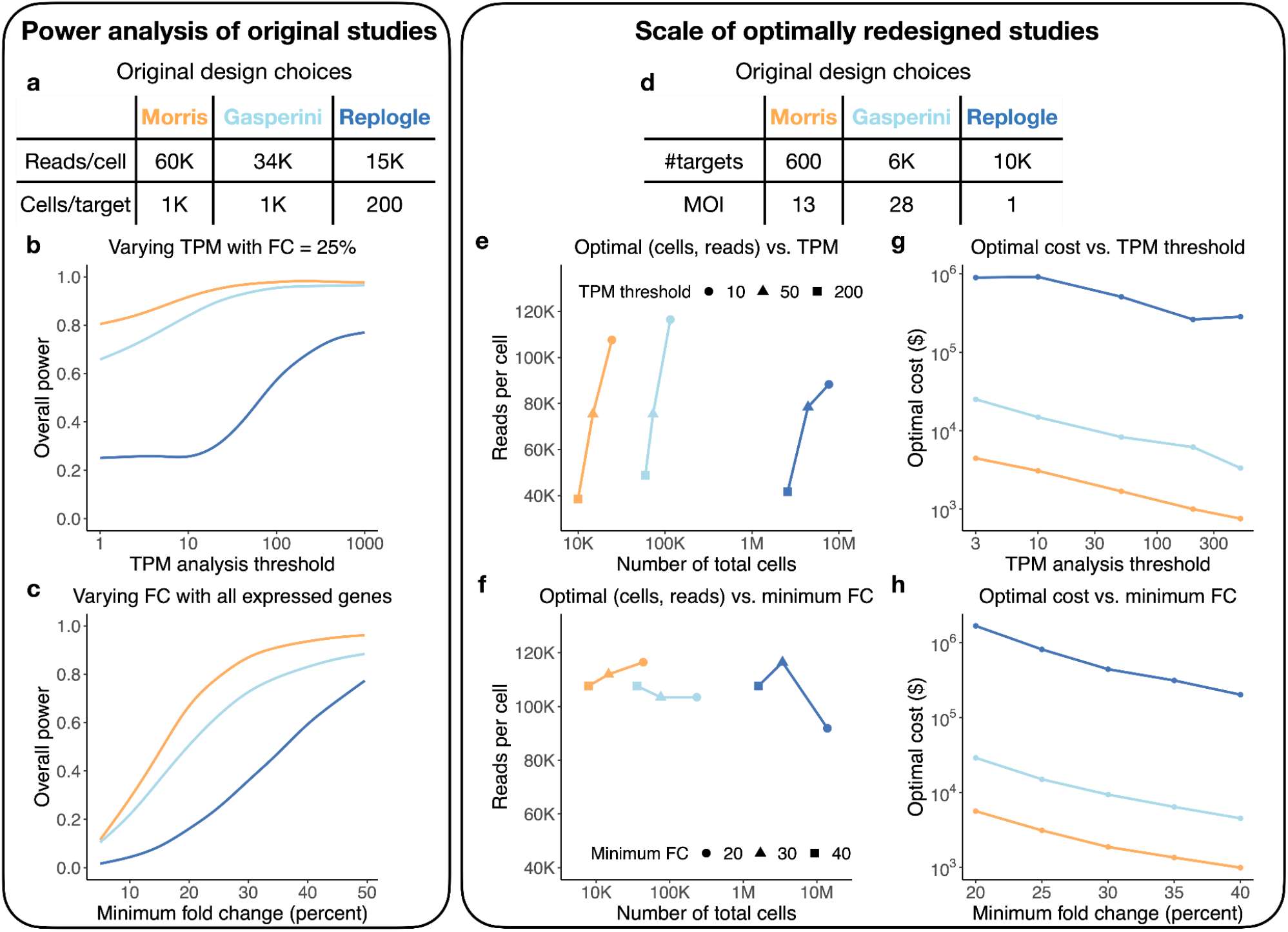
PerturbPlan unveils differences in power and cost across three Perturb-seq studies. **a, d:** Original design information across three Perturb-seq studies.^7,8,11^ **b-c:** Power comparison against different TPM analysis thresholds and FCs of interest. The power is computed with cells per target and reads per cell in panel **a**. **e-f**: Optimal reads per cell and number of total cells across TPM thresholds and FCs. The power of interest is 0.8 and the design choices in panel **d** are used for design optimization. 25% FC is considered when varying TPM threshold (**e**) and genes with at least 10 TPM are considered when varying FC (**f**). **g-h**: Optimal cost against different TPM analysis thresholds and effect sizes of interest while varying reads per cell and cells per target. Details the same as in panels **e**-**f**.

We then used PerturbPlan to explore prospective redesigns of these experiments. Specifically, we fixed the target power at 0.8 and optimized total cost over reads per cell and cells per target for given combinations of TPM threshold and effect size, using the scale information in the original designs (**Fig. 4d**) and 2025 prices for library preparation and sequencing (**Methods**). Comparing across the three studies, we found that achieving 80% power for any TPM threshold and FC requires roughly the same number of reads per cell but more total cells for Replogle than for Gasperini, and for Gasperini than for Morris (**Fig. 4e,f**). For example, to detect a 20% FC, the Morris, Gasperini, and Replogle redesigns required 43K, 233K, and 13.9M cells, respectively (**Fig. 4f**), translating to costs of $5.6K, $29K, and $1.7M (**Fig. 4h**). One conclusion is that obtaining good power for a low-MOI screen targeting all expressed genes is very expensive (at least, with our cost estimates). We also found that changing the TPM threshold and FC had differential impacts on the optimal designs. In particular, we found that changing the TPM threshold had a significant impact on the optimal reads per cell while changing the FC did not, instead primarily impacting the optimal numbers of cells. This observation is likely due to the fact that decreasing the TPM threshold includes more lowly expressed genes, which in turn requires more reads per cell to obtain sufficient UMIs per gene to achieve the target power. On the other hand, this phenomenon does not occur when changing FC. Finally, our results quantified the cost of achieving a more informative experiment, either via including more genes by lowering the TPM threshold (**Fig. 4g**) or having power to detect weaker regulatory relationships by lowering the minimum FC (**Fig. 4h**). For example, for the Morris study, decreasing the TPM threshold from 500 to 3 or decreasing the FC from 40% to 20% require approximately a ten-fold increase in experimental cost.

### Power and cost comparison between Perturb-seq and TAP-seq

Targeted Perturb-seq^6^ (TAP-seq) enables efficient expression profiling in single cells by focusing sequencing depth on genes of interest. Using two rounds of semi-nested multiplex PCRs with gene-specific primers, TAP-seq amplifies and then sequences transcripts from a prespecified set of genes rather than the whole transcriptome. However, poorly performing TAP-seq primers lower the PCR amplification efficiency and can negatively impact power for their target genes.^6^ We applied PerturbPlan to quantify the impacts of these differences on power and cost between TAP-seq and Perturb-seq platforms. For this analysis, we drew on Perturb-seq data from Gasperini et al.^7^ and TAP-seq data from Ray et al.,^9^ both in K562 cells. The Ray dataset targeted 303 genes, which in our analysis we consider the genes of interest. We also use UMIs per cell at saturation as a depth-agnostic measure of gene expression that is appropriate for Perturb-seq and TAP-seq (**Methods**).

As a starting point, we used PerturbPlan to fit saturation curves to both datasets, restricting to the set of genes of interest (**Step 2** in **Fig. 2** and **Methods**). We confirmed that much fewer sequenced reads per cell are required to obtain the same number of UMIs per cell across targeted genes, as a result of a much higher fraction of reads mapping to these genes (52.7% versus 3.7%, **Fig. 5a**). On the other hand, we observed poor efficacy for some TAP-seq primers, leading to lower numbers of UMIs per cell at saturation for those genes in TAP-seq compared to Perturb-seq (**Fig. 5b**). Otherwise, the UMIs per cell at saturation across genes agree reasonably well across the two datasets (**Fig. 5b**). It is sensible to introduce an expression threshold in constructing the perturbation-gene pairs to analyze in a TAP-seq dataset, in order to prevent genes with poor primers from impacting power. We used PerturbPlan to calculate the power for testing these filtered perturbation–gene pairs under two scenarios—profiling only targeted genes versus profiling the whole transcriptome—assuming a 10% FC (**Fig. 5c**). These scenarios correspond to TAP-seq and Perturb-seq, respectively. We carried out a similar analysis, this time varying the FC while fixing the expression threshold to 1 UMI per cell (**Fig. 5d**). We found that at every expression threshold and FC tested, TAP-seq achieved a higher power than Perturb-seq.

**Figure 5.**
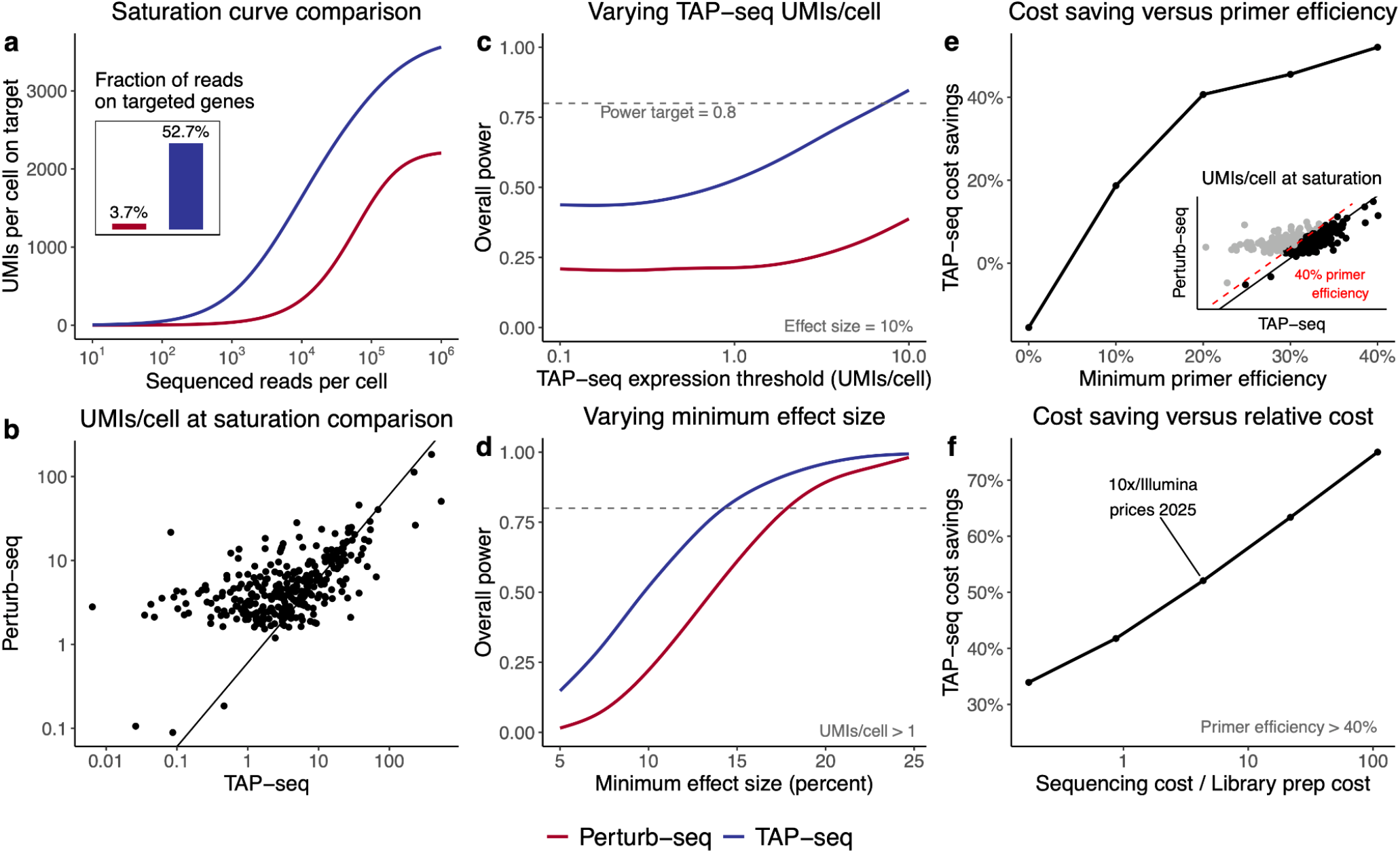
PerturbPlan quantifies the advantages of TAP-seq over Perturb-seq and their limits. **a:** Sequencing saturation curve comparison between TAP-seq and Perturb-seq. The inset plot shows the fraction of reads mapped to the targeted genes in the two assays. **b:** UMIs per cell at saturation comparison between Perturb-seq and TAP-seq. UMIs per cell at saturation are computed using PerturbPlan’s saturation curve fitting (**Fig. 2, Step 2**). The diagonal line represents the expected UMIs per cell at saturation for each gene under a scenario where the relative expressions of all targeted genes are the same across Perturb-seq and TAP-seq. **c-d:** Power comparison across TAP-seq expression thresholds and effect sizes. We filter genes based on the UMIs per cell at saturation obtained from TAP-seq data. The power is computed with cells per target and reads per cell from the TAP-seq study.^9^ **e:** TAP-seq cost savings against primer efficiency threshold when the desired power is 80%. The primer efficiency is thresholded based on the ratio of UMIs per cell measured in TAP-seq to that measured in Perturb-seq. When the minimum primer efficiency is 0, all the genes are preserved in the analysis set. The inset plot illustrates the filtered genes (black points) passing a 40% primer efficiency threshold. **f:** TAP-seq cost savings against relative cost of sequencing and library preparation when the desired power is 80%. Sequencing cost is defined as dollar per million raw reads and library preparation cost is defined as dollar per captured cell.

We then turned to quantifying the cost savings offered by TAP-seq, compared to a hypothetical Perturb-seq experiment involving the same perturbations. To this end, we used PerturbPlan to design the optimal reads per cell and cells per target to achieve 80% power while holding other experimental parameters fixed (e.g., number of targets, number of gRNAs per target). Given the optimized reads per cell and cells per target, we compared the total costs of the optimal TAP-seq and Perturb-seq experiments. In particular, we examined how primer efficiency and the relative costs of sequencing and library preparation influence the advantage of TAP-seq (**Fig. 5e,f**). We defined primer efficiency as a scaled ratio of UMIs per cell at saturation in TAP-seq versus Perturb-seq for a given gene (**Methods**). As an illustration, genes with at least 40% primer efficiency are highlighted in black in the inset of **Fig. 5e**. When the minimum primer efficiency threshold is set to 0, all targeted genes from the Ray study are retained, including those with substantially lower expression than in Perturb-seq (**Fig. 5b**). In this regime (left-most point in **Fig. 5e**), TAP-seq becomes disadvantageous, driven by poor primer performance for lowly expressed genes. As the primer-efficiency threshold is raised and poorly performing primers are excluded, TAP-seq becomes increasingly favorable. Focusing on genes with high primer efficiency (**Fig. 5f**), the cost savings from TAP-seq grow as the ratio of sequencing cost (dollars per million reads) to library preparation cost (dollars per captured cell) increases. This is intuitive: the primary advantage of TAP-seq over Perturb-seq is the reduced reads-per-cell requirement to achieve a given number of UMIs per gene, so its relative benefit is amplified when sequencing is relatively more expensive than library preparation.

In sum, PerturbPlan revealed that while TAP-seq often offers an advantage over Perturb-seq in terms of power and cost, the extent of this advantage depends crucially on the efficacy of the TAP-seq primers and the relative costs of sequencing and library preparation (which vary as both technologies evolve). In some cases, the poor primer design can eliminate TAP-seq’s advantage entirely. When applied for experimental design, PerturbPlan can help experimentalists quantify and navigate these trade-offs. Beyond designing optimal TAP-seq or Perturb-seq experiments, PerturbPlan can help guide decisions such as whether to redesign the primer panel to improve efficiency or even whether to choose TAP-seq versus Perturb-seq for a given study.

## Discussion

PerturbPlan facilitates versatile, fast, and interactive single-cell CRISPR screen experimental design. Compared with the only other existing power analysis tool for this assay,^9^ PerturbPlan reduces the sequential runtime to sweep across 1000 design configurations from 3.8 months to 1 second. This enables a paradigm shift in single-cell CRISPR screen experimental design from submitting jobs on high-performance clusters to interacting with a web application, at a crucial time when these assays grow in both scale and adoption. Compared to the existing scPower tool^24^ for single-cell differential expression, PerturbPlan provides a fully tailored solution for single-cell CRISPR screen design.

PerturbPlan has a few limitations, which we look forward to addressing in future work. Currently, PerturbPlan allows for the specification of effect sizes in terms of a single FC of interest, and an expected proportion of tested perturbation-gene pairs with at least that FC. This allows for conservative experimental design, but does not fully account for effect size heterogeneity. Furthermore, PerturbPlan does not have functionality to estimate this non-null proportion from pilot data. Finally, PerturbPlan does not currently accommodate covariates beyond library size, such as batch.

While PerturbPlan is a tool for prospective power analysis and experimental design, it is sometimes of interest to analyze power retrospectively, i.e., after the data are collected. In particular, retrospective power analyses are helpful for curating perturbation-gene associations (and non-associations) for predictive model training.^13^ In such analyses, more information is available for power calculation, but power is desired on a per-perturbation-gene-pair basis rather than on average across pairs (the latter is what PerturbPlan provides). We believe that some of PerturbPlan’s components can be repurposed for such retrospective analyses, but this is beyond the scope of the current work.

PerturbPlan’s experimental design features are complementary to those of recent tools leveraging perturbation effect prediction for nominating genomic elements to target in single-cell CRISPR screens.^25–27^ A promising direction would be to integrate PerturbPlan with such tools, in order to account for power, cost, and predicted perturbation effects in a unified framework. Using the genes predicted to be impacted by the chosen perturbations could also help guide the choices of genes to capture within TAP-seq.

In conclusion, we expect that PerturbPlan will accelerate the discovery of regulatory relationships by making it easier for researchers to design powerful and cost-effective single-cell CRISPR screens.

## Methods

### PerturbPlan power formula

For clarity, we present here a version of the power formula used for left-sided tests (right- and two-sided tests are analogous), without incorporating additional accelerations. A more comprehensive exposition of the power formula (including accelerations), as well as its derivation, is presented in the **Supplementary Note**.

#### Inputs

Reference expression data:

- UMI counts *Y*_*ij*_ for cells *i*and genes *j*
- Number of mapped reads *R*_*u*_ for each observed UMI *u* =1, …,*U*
- Number of cells *N*_*ref*_
- Mapping efficiency *e*∈ [0, 1]

Desired effect sizes:

- Minimum FC of interest µ
- gRNA variability σ^2^ ≥ 0
- Minimum proportion of non-null pairs π ∈ [0, 1]

Experimental choices:

- Sequenced reads per cell *R*_*seq*_
- Number of cells *N*
- Number of targeted elements *L*
- Number of gRNAs per target *K*
- Number of non-targeting gRNAs *K*_0_
- Multiplicity of infection MOI

Analysis parameters:

- FDR target level *q*
- Test sidedness: left, right, or two-sided
- Element-gene pairs of interest {(*l,j*) : (*l,j*) ∈ 𝒫}
- Control group: non-targeting cells or complement cells

#### Step 1: Baseline expression summarization

Fit a negative binomial model to the counts {*Y*_*ij*_}_*i*_ of each gene *j* across cells to obtain maximum likelihood estimates of α_*j*_ (relative expression of gene *j*, summing to 1 across *j*) and θ_*j*_ (dispersion of gene *j*):

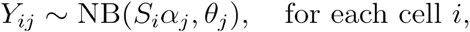

where *S*_*i*_ is the total number of UMIs in cell *i*.

#### Step 2: Library size computation

Using the read data {*R*_*u*_}_*u*_ and the `preseqR.rSAC()` function from the `preseqR` package, fit a function 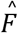 mapping the total number of mapped reads to the total number of UMIs represented among these reads. For *R*_*seq*_ planned sequenced reads per cell, compute the For average library size per cell via

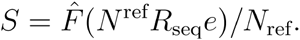

This formula involves rescaling the function 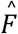 from `preseqR` to the per-cell level and translating sequenced reads to mapped reads via the mapping efficiency *e*.

#### Step 3: Test statistic distributional approximation

For each (*l, j*) ∈ *P*, carry out the following steps to get a normal approximation *T*_*lj*_ ~ *N*(η_*lj*_, τ_*lj*_). First, independently sample per-gRNA effect sizes β_*kl,j*_ ~ *N*(µ, σ^2^), and define 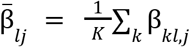. Next, compute the expected number of “treatment” cells *N*^*t*^ bearing a perturbation of element *l* and the expected number of “control” cells *N*^*c*^ used for comparison:

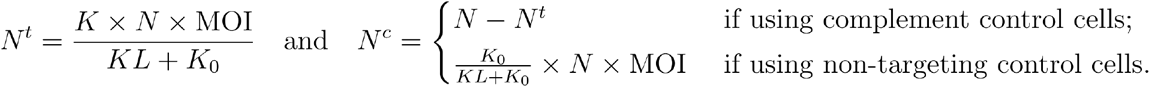

Then, compute

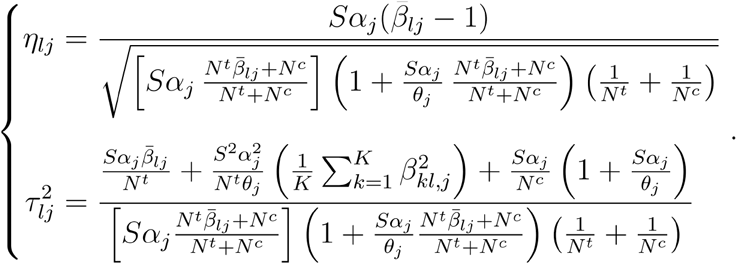

#### Step 4: BH cutoff approximation

Use a bisection algorithm to find the solution to

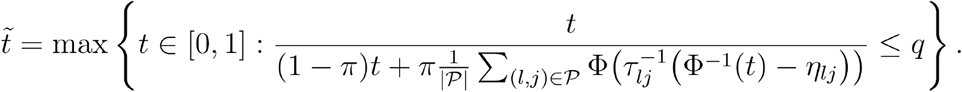

#### Step 5: Overall power calculation

Compute the overall power via

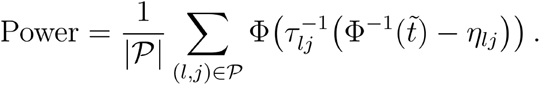

### PerturbPlan experimental design optimization

PerturbPlan uses a simple and flexible cost model for single-cell CRISPR screens, requiring users to specify a cost per cell and a cost per read. This parameterization of cost accommodates for a diversity of experimental platforms. Given user-specified cost parameters, we use grid search to optimize experimental designs. To improve computational efficiency, we combine grid search with adaptive search strategies. We categorize the grid search strategies for the design problems listed in **Extended Data Tab. 1** into three types. Category (a) comprises designs that optimize a single non-cost parameter subject only to a lower bound on power; these are designs 1-4. Category (b) comprises designs that minimize total cost while jointly varying cells per target and reads per cell under a power constraint; this is design 5. Category (c) comprises designs that minimize either the detectable fold change or the expression threshold subject to both a power constraint and a budget constraint, while allowing one or both profiling-depth variables (cells per target and reads per cell) to vary; these are designs 6-11. Within category (c), designs 6 and 9 vary both profiling-depth variables, whereas designs 7-8 and 10-11 vary only one.

Below, we describe the adaptive grid search strategy used for each category. Let *Pow* denote the target power specified by the user.

#### Design category (a)

When optimizing a single non-cost parameter, we evaluate power on a grid of 20 points. For effect size or expression threshold, where power is more sensitive, we use prespecified grids of 15 points. Specifically, for the expression threshold, we use equally spaced quantiles of the TPM values of genes in the reference expression dataset. For effect size, we use equally spaced values in [0.5, 0.9] for knockdown effects, [1. 1, 1. 5] for activation effects, or the union of both intervals when both effects are of interest. For reads per cell and cells per target, which can vary substantially across experimental designs, we adopt different search strategies. For reads per cell, we use a heuristic range based on the saturation curve: from 0.1 times to 5 times the estimated UMI count at saturation, and evaluate power at 20 logarithmically spaced values within this range. For cells per target, we use an adaptive two-step procedure. First, we identify the minimum and maximum values of the parameter that achieve power levels of *max*{*Pow* − 0. 3, 0. 05} and *min*{*Pow* + 0. 3, 0. 95},respectively. We then evaluate power at 20 logarithmically spaced values between these bounds.

#### Design category (b)

When optimizing cost, both reads per cell and cells per target are varied, requiring a more refined grid search. For reads per cell, we use the same heuristic range as in design category (a), spanning from 0.1 times to 5 times the estimated UMI count at saturation. For cells per target, we set the minimum value as the number of cells that, together with the maximum reads per cell, yields power *max*{*Pow* − 0. 3, 0. 05}, and the maximum value as the number of cells that, together with the minimum reads per cell, yields power *min*{*Pow* + 0. 3, 0. 95}. We then evaluate power on a grid of 20 logarithmically spaced values along each dimension, resulting in 400 (cells per target, reads per cell) combinations.

#### Design category (c)

This category is the most computationally intensive because it involves joint optimization over three variables. To reduce computational cost, we first select five equally spaced values for the expression threshold or effect size using the same strategy as in category (a). For each fixed value, we then apply the adaptive grid search described in category (b) to identify combinations of cells per target and reads per cell that achieve the target power Pow. Finally, we compute the cost for each feasible combination and select the optimal design subject to the cost constraint.

### Software implementation

The PerturbPlan web application is implemented as a Shiny front end distributed in the open-source `perturbplanApp` R package. The application can be run from a browser via perturbplan.com or locally (e.g., for analyses on sensitive data) by downloading `perturbplanApp` and calling the run_app() function. `perturbplanApp` depends on a separate open-source R package, `perturbplan`, which implements all power calculations, optimization routines, and utilities for processing user-provided custom reference expression data. The core power calculations are implemented in Rcpp.

### Simulation validation details

We use the simulation setup proposed in the power analysis pipeline,^9^ which is available at: https://github.com/ZiangNiu6/Sceptre_Power_Simulations. The simulation pipeline estimates power for a given parameter configuration as follows: (1) apply `sceptre`^28,29^ to the sampled perturbation and gene expression matrices (with the sampling procedure described below); (2) combine the resulting `sceptre` p-values with additional null p-values drawn from the uniform distribution, and apply the Benjamini–Hochberg procedure at a target FDR level of 0.1 to determine the set of significant pairs; and (3) repeat steps (1)–(2) 200 times and define power as the average fraction of non-null pairs that are declared significant. We consider a total of 7,364 pairs involving 526 distinct genes. The number of cells per target ranges from 250 to 2,000 in increments of 250, and effect sizes range from 5% to 40% in increments of 5%. When varying the effect size, we fix the number of cells per target at 1,000; conversely, when varying the number of cells per target, we fix the effect size at 15%. Throughout, we set the non-null proportion to 0.05. The details of data generating procedure can be found in the section “Simulation benchmarking details” of **Supplementary Note**. For each of the 16 parameter configurations, we ran the simulation pipeline in parallel on a cluster and summed the runtimes across processors to obtain CPU-hours. Jobs were executed on a virtualized x86_64 compute node running Rocky Linux 8, equipped with an Intel Xeon Platinum 8375C CPU (2.9 GHz).

### Real data validation details

#### Preprocessing 10x, Gasperini and Ray datasets

We analyze three K562 datasets for real-data validation: two Perturb-seq studies (the 10x dataset and the Gasperini dataset) and one DC-TAP-seq study (the Ray dataset; see the “Data Availability” section). We process the FASTQ files using Cell Ranger 8.0.1 and work primarily with two outputs: molecule_info.h5, which contains the mapped-read information, and filtered_feature_bc_matrix, which contains the UMI count data. The 10x and Gasperini datasets are used to validate the saturation-curve fitting method, whereas the Gasperini and Ray datasets are used to evaluate the power of PerturbPlan. For the latter two datasets, we define positive and negative controls as follows. As positive controls, we include known enhancer-gene pairs and perturbations targeting transcription start sites (TSSs) of genes in the Gasperini dataset, as well as TSS-targeting perturbations in the Ray dataset. As negative controls, we use non-targeting gRNAs.

#### Downsampling validation for saturation curve fitting

The saturation curve modeling workflow is validated using the `molecule_info.h5` data from the 10x and Gasperini datasets. We first fit the preseqR model on the Gasperini dataset by downsampling sequencing reads from 1% to 100%, pairing each downsampled dataset with the corresponding full-depth data (100% reads), and selecting the parameter estimates from the downsampling–full-depth pair with the lowest training loss. The dataset contains 32 SRRs distributed across two runs, and we use either 1 or 8 SRRs from one of the runs to estimate the model parameters: the total number of UMIs per cell and the variation parameter. We then validate the fitted saturation curve on the downsampled 10x dataset (**Fig. 3d**). We additionally benchmark our approach against a competing method based on the `umi.read.relation()` function in `scPower` (**Extended Data Fig. 3a-b**). In the competing method, UMI abundance is predicted from sequencing depth by jointly fitting a linear regression of UMI counts on the log-transformed total read counts across four read-downsampled datasets with sampling ratios of 25%, 50%, 75%, and 100%. To assess robustness of our method with respect to sequencing depth, we examine the fit quality under different downsampling choices (**Extended Data Fig. 3c,d**).

#### Downsampling workflow for power validation

We begin by fixing a combination of the number of cells (*N*) and reads per cell (*R*); the procedure for choosing these parameters is described in the next paragraph. Our validation then proceeds in four steps. (1) Starting from the `molecule_info.h5` file (after removing background barcodes), we generate 200 independent downsampling replicates, each with *N* cells and *R* reads per cell. (2) For each replicate, we construct the subsampled gene expression matrix and subset the gRNA perturbation matrix using the cell barcodes present in the gene expression matrix. (3) We then run `sceptre` using the resulting gene and gRNA expression matrices to obtain p-values for each of the 200 downsampling replicates. (4) Finally, we apply the Benjamini-Hochberg procedure to control for multiplicity, define the discovery set, and compute power as the average number of detected positive-control pairs across the 200 replicates. A key determinant of power in this setting is the proportion of non-null element-gene pairs. For validation, we include only negative-control and positive-control pairs (both defined as in the “Preprocessing 10x, Gasperini and Ray datasets” section), so that the non-null proportion is fixed by design. We set the non-null proportion to 0.01 when downsampling the Ray data and to 0.1 when downsampling the Gasperini data.

For the Gasperini dataset, we let *R* take 10 values on an equally spaced grid between 4,423 and 22,117, and *N* take 10 values on an equally spaced grid between 18,944 and 75,792. When varying *R*, we fix *N* =37, 888; when varying *N*, we fix *R* =13, 270. We filter the positive controls to those with effect sizes between 25% and 30% for low effect-size variability, and between 25% and 40% for high effect-size variability. For the Ray dataset, we let *R* take 10 values on a logarithmically spaced grid between 255 and 25,473, and *N* take 10 values on an equally spaced grid between 15,280 and 84,050. When varying *R*, we fix *N* =76, 410; when varying *N*, we fix *R* =25, 473. In addition, for the Ray dataset we downsample the gRNAs so that each target is represented by 10 gRNAs out of the original 15. We filter the positive controls to those with effect sizes between 15% and 20% for low effect-size variability, and between 15% and 30% for high effect-size variability.

#### UMIs per cell at saturation and primer efficiency

To compute a gene’s UMIs per cell at saturation for either Perturb-seq or TAP-seq, we extracted the total UMIs per cell at saturation from the fitted saturation curve, and then multiplied by the gene’s relative expression. To define the efficiency of a primer, our starting point was the ratio of TAP-seq UMIs per cell at saturation to Perturb-seq UMIs per cell at saturation. However, TAP-seq recovered more total UMIs per cell at saturation across targeted genes than Perturb-seq (**Fig. 5a**), potentially due to the differences in total transcriptional activity across the two K562 datasets analyzed. For a fair comparison, we therefore defined primer efficiency for a gene via

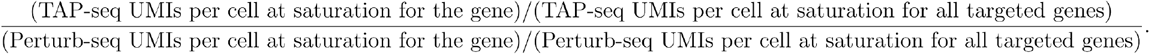

This definition is framed in terms of relative gene expression, and has the property that decreasing one gene’s primer efficiency leads to an increase in other genes’ primer efficiencies. In this sense, our definition of primer efficiency is fundamentally relative to those of other primers, and it is possible to have primers with relative efficiencies greater than 1.

### Additional design parameters in case studies

Across all case studies, we set the non-null proportion to 0.005 and target an FDR level of 0.1, using a left-sided test for knockdown effects. For the default effect size specification, we fix the gRNA variability parameter σ (introduced in “PerturbPlan power formula” section) at 0.13, following a previous work.^13^

#### Experimental cost parameters

We used **$0.086 per captured cell** and **$0.374 per million sequenced reads**, as obtained from the 2025 costs of 10x Genomics library preparation followed by Illumina sequencing. In particular, for library preparation, we considered the 10x Genomics Chromium Next GEM Single Cell 3′ Gene Expression v4 workflow with direct capture of perturbation identities using the 10x Genomics Feature Barcode technology (guide/feature barcode capture). Library-preparation consumables for a full 16-lane experiment totaled $27,510, comprising the Chromium Next GEM Single Cell 3′ Reagent Kits v4 ($23,670), Chromium Next GEM Chip(s) ($1,250), Feature Barcode reagents / guide capture kit ($670), and Dual Index Kit / indexing reagents ($1,920). With an expected recovery of approximately 320,000 cells across 16 lanes, this corresponds to an estimated library-prep cost of $0.086 per recovered cell. Sequencing costs were estimated using an Illumina NovaSeq X Series 25B (100-cycle) kit priced at $9,360, corresponding to approximately $0.374 per million reads based on nominal kit output.

#### Perturb-seq study comparisons

We set the number of gRNAs per element based on the original studies: Morris study used 3 gRNAs per element, Gasperini study used 2, and Replogle study used 1. For all three assays, we assume a mapping efficiency of 0.704, defined as the ratio of mapped reads per cell to raw reads per cell (mapped/raw). This value is estimated by dividing the number of mapped reads per cell by the number of raw reads per cell and has been observed to be relatively stable across Perturb-seq studies, so we apply the same value to all datasets.

#### TAP-seq and Perturb-seq study comparisons

We set the number of gRNAs per element to 15 in the Ray study and 2 in the Gasperini study, matching the values used in the original publications. For mapping efficiency, we use 0.704 for the Gasperini study and 0.371 for the Ray study. The mapping efficiencies are obtained from the Cell Ranger summary outputs.

## Data Availability

- Source data for all figures: **Supplementary Table 1**.
- 10x K562 dataset: https://www.10xgenomics.com/datasets/10k-k562-transduced-with-small-guide-library-5-ht-v2-0-chromium-x-2-standard
- Gasperini K562 dataset: https://www.ncbi.nlm.nih.gov/geo/query/acc.cgi?acc=GSE120861
- Ray K562 dataset: https://data.igvf.org/analysis-sets/IGVFDS7288SJVF/

## Code Availability

- Web application: perturbplan.com
- Web application documentation: https://katsevich-lab.github.io/perturbplanApp
- Web application source code: https://github.com/Katsevich-Lab/perturbplanApp
- Computational backend and data preprocessing R package documentation: https://katsevich-lab.github.io/perturbplan/
- Computational backend and data preprocessing R package source code: https://github.com/Katsevich-Lab/perturbplan
- Code to replicate all analyses: https://github.com/Katsevich-Lab/perturbplan-replication

## Acknowledgements

E.K. acknowledges support from NSF DMS-2310654. We thank the Wharton Research Computing team for their computing support, and Stanford University and the Stanford Research Computing Center for providing computational resources and support as part of the Sherlock High-Performance Compute Cluster. J.E. acknowledges support from the NIH-NHGRI IGVF Consortium (UM1HG011972), the NIH-NHGRI MorPhiC Consortium (U01 HG013176), NIH-NHLBI (P01HL180323), the Applebaum Foundation, and the Novo Nordisk Foundation Center for Genomic Mechanisms of Disease (Novo Nordisk Fonden NNF21SA0072102). L.M.S and A.R.G. acknowledge support from the National Human Genome Research Institute of the National Institutes of Health (RO1HG011664 and the IGVF consortium via UM1HG011972). We thank members of the IGVF Consortium for their feedback on this work.

## Author Contributions

E.K. conceived the study. E.K and J.M.E. supervised the project. Z.N. and Y.H. implemented the PerturbPlan web application and R package, and carried out the empirical studies, with assistance and guidance from A.R.G., J.G., and J.M.E. J.G. and A.R.G. developed and compared the simulation-based power analysis pipeline with input from L.M.S. and J.M.E. J.R. contributed DC-TAP-seq data and advised on experimental parameters. E.K., Z.N., and Y.H. wrote the manuscript, with input from A.R,G. and J.M.E.

## Extended Data

### Extended Data Figures

**Extended Data Figure 1.**
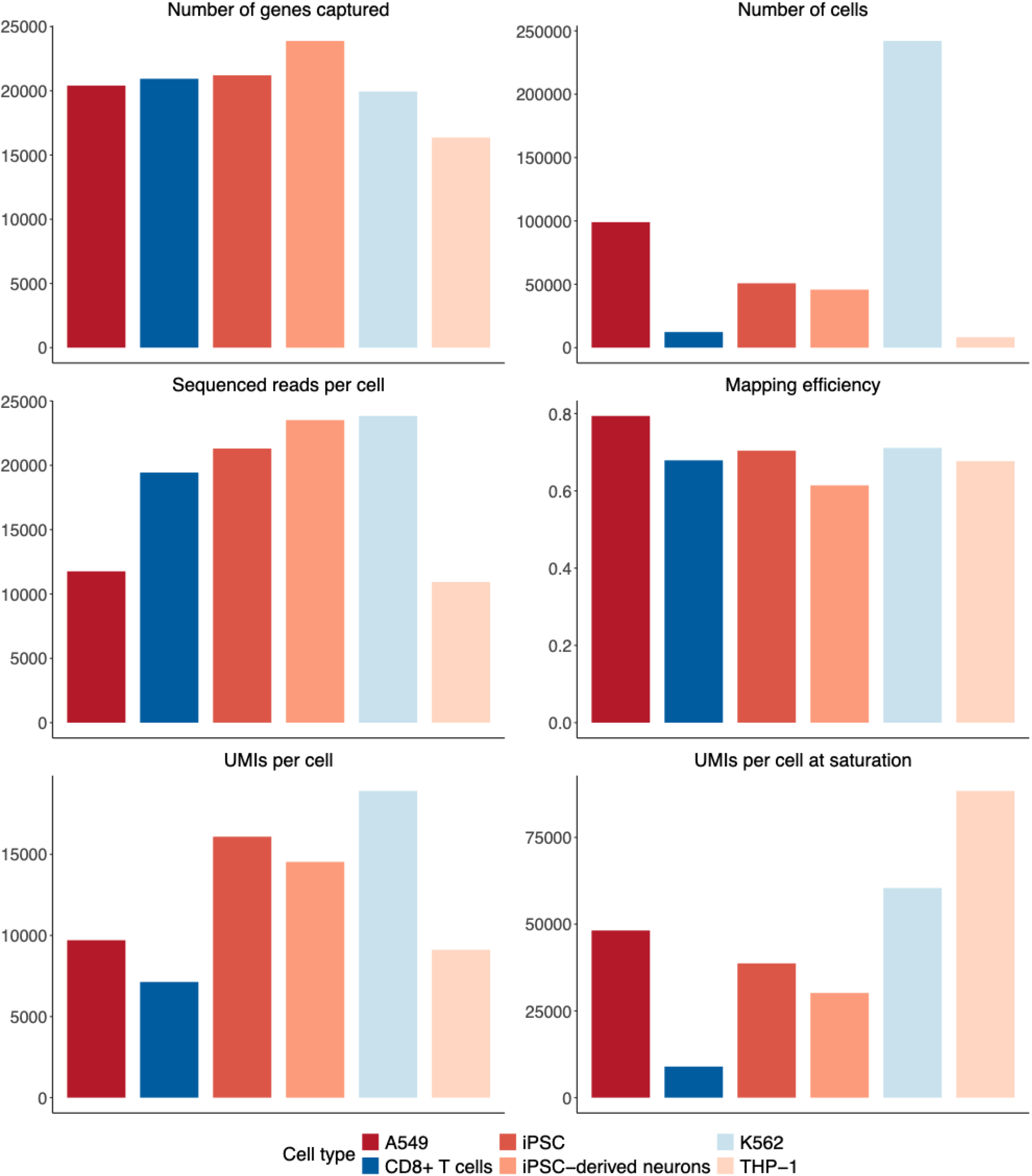
Properties of the 6 reference expression datasets built into PerturbPlan. **Number of genes captured:** Total number of distinct genes detected in each dataset. **Number of cells:** Number of cells in reference expression data. Note that only cells carrying non-targeting gRNAs were used for the THP-1 cell type, while all cells were used for the other cell types. **Sequenced reads per cell:** Average number of sequencing reads confidently mapped to transcriptome per cell. **Mapping efficiency:** Fraction of reads that are confidently mapped to the transcriptome. **UMIs per cell:** Average UMIs detected per cell. **UMIs per cell at saturation:** Estimated UMIs per cell at sequencing saturation based on library complexity modeling.

**Extended Data Figure 2.**
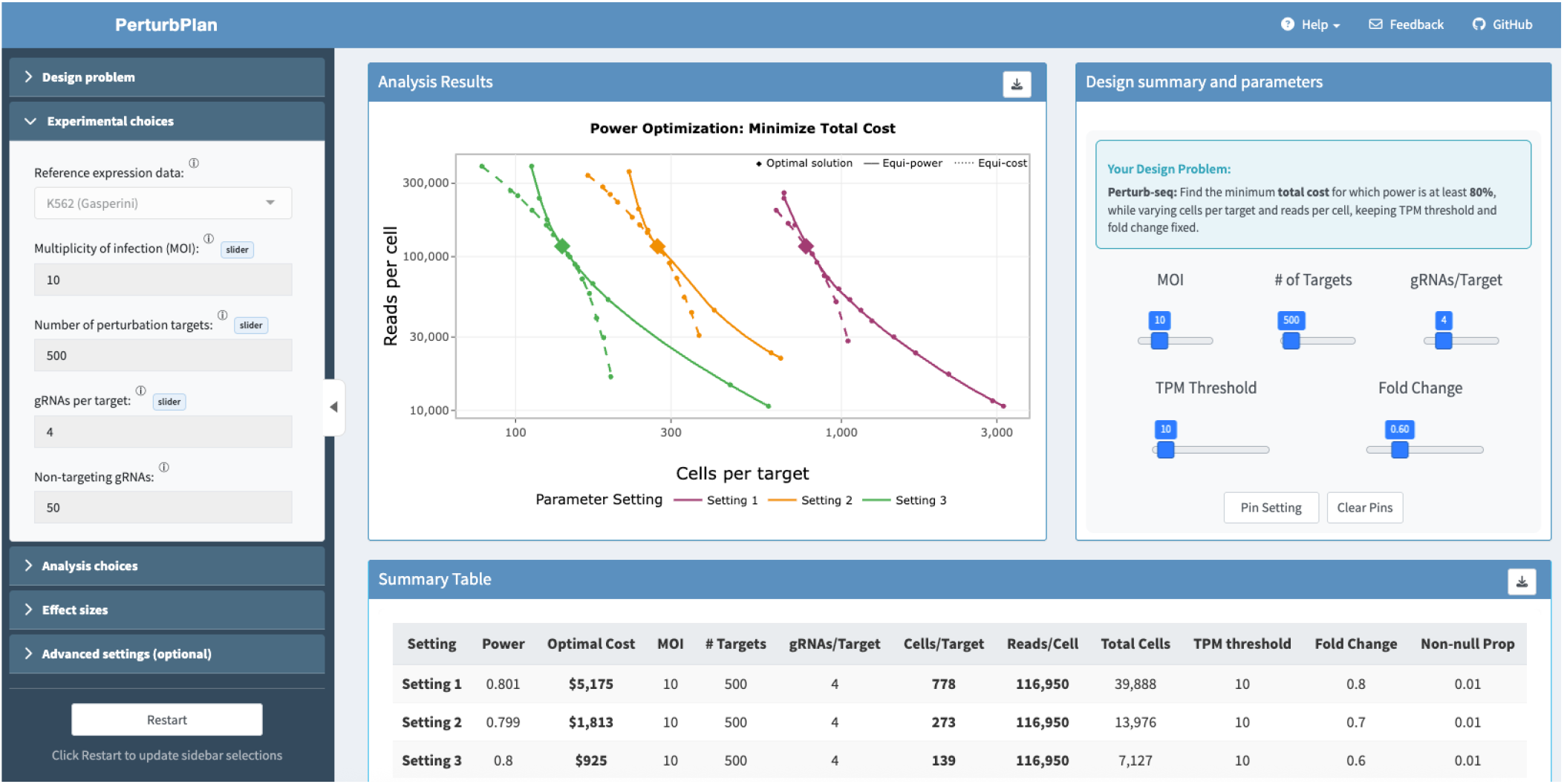
Screenshot of PerturbPlan web application. **Sidebar:** The sidebar at left facilitates initial user input of design problem, experimental choices, analysis choices, effect sizes, and advanced settings (optional). The sidebar has a button initially labeled “Plan”, which changes to “Restart” upon clicking. **Analysis results:** A graphical display of the experimental design optimization for each user-chosen setting, including a button in the top-right corner to download the plot in PDF format. **Design summary and parameters:** A text summary of the user’s design problem, as well as sliders to adjust parameter settings. At the bottom, buttons are available to pin the current setting (which fixes the corresponding curve in “Analysis results” and adds a legend entry) as well as to clear the pins. **Summary table:** A tabulation of the parameter settings pinned by the user, the optimal designs suggested by PerturbPlan (in bold), and the resulting power and cost. A button in the top-right corner allows downloading this table and additional supporting results in Excel format. **Navigation bar:** Across the top, a “Help” button providing links to consult for documentation, questions and feature requests, and bug reports; a “Feedback” button to engage privately with the developers, and a “GitHub” button to be taken to the source code.

**Extended Data Figure 3.**
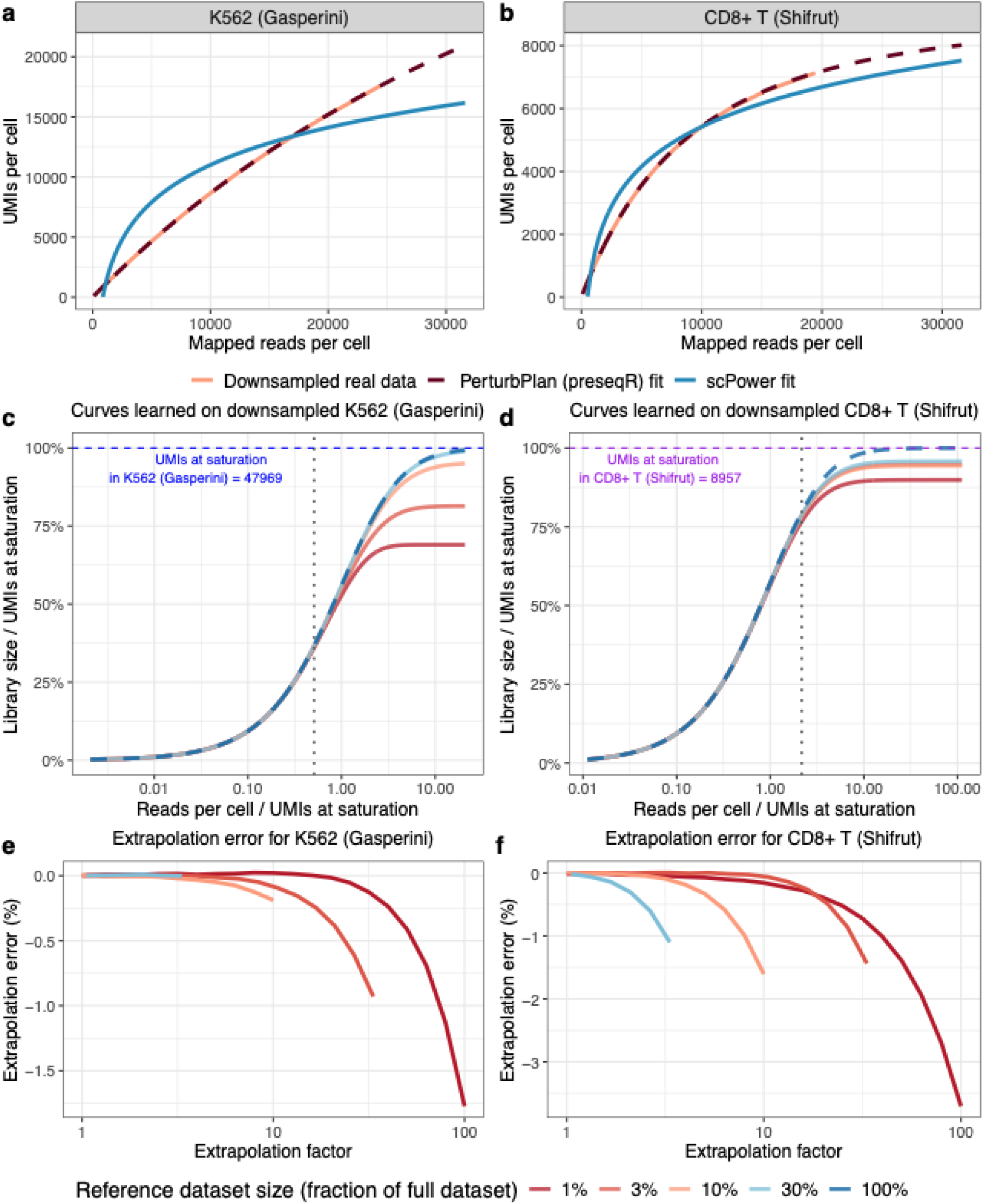
PerturbPlan’s preseqR-based model fits sequencing saturation curves accurately across a range of sequencing depths of reference datasets. **a-b:** Sequencing saturation curves learned by scPower and PerturbPlan against the ground truth for two different reference datasets: Gasperini K562 (Gasperini), a shallowly sequenced reference dataset and CD8+ T cells (Shifrut), a deeply sequenced reference dataset (**b**). **c–d:** Sequencing saturation curves learned from downsampled subsets of the reference datasets. Curves are learned on K562 (Gasperini) data **(c)** and CD8+ T cells (Shifrut) data **(d)** using subsets with total read counts ranging from 1% to 100% of the full reference data. In each panel, the dashed horizontal line indicates the UMIs per cell at saturation estimated from the full dataset, and the dashed vertical line indicates the sequencing depth of the full reference dataset. The dashed curve is identical to the PerturbPlan (preseqR) fit shown in panel a, after rescaling of the axes. **e–f:** Extrapolation error as a function of extrapolation factor for K562 (Gasperini) **(e)** and CD8+ T cells (Shifrut) **(f)**. Errors are computed by comparing extrapolated UMIs at saturation from downsampled pilot datasets (1%–100% of the full dataset) to the saturation level inferred from the full reference dataset. The extrapolation factor is defined as the ratio between the sequencing depth at the estimation point and the sequencing depth of the downsampled dataset used for training.

**Extended Data Figure 4.**
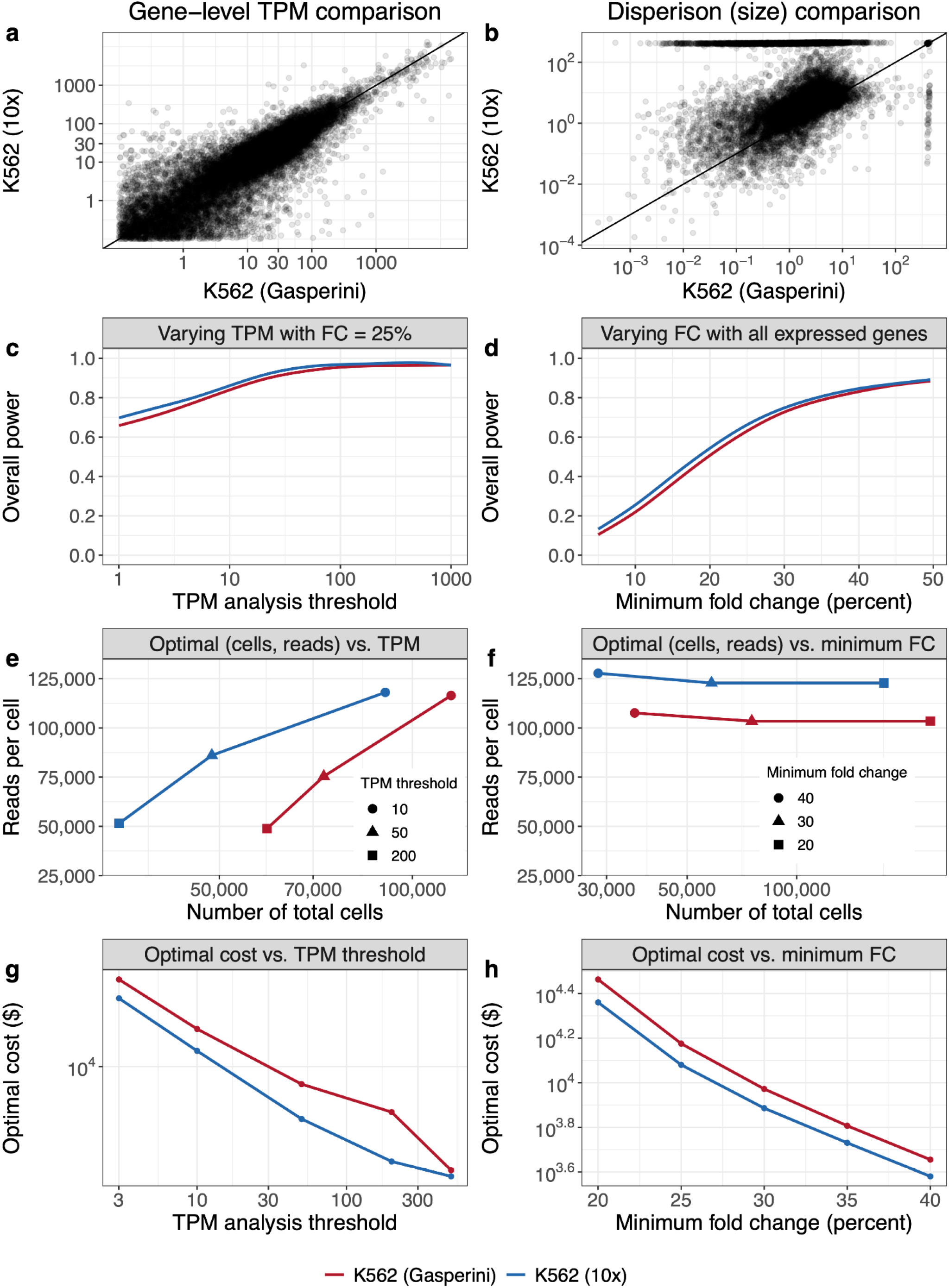
PerturbPlan results are robust to different reference data from the same cell type. **a-b:** Comparison of relative expression and dispersion (size parameter) across two Perturb-seq experiments in the same cell type. **c-d:** Power comparison against different TPM analysis thresholds and FCs of interest calculated from reference expression datasets in K562. Similar to the setup in **Fig. 4b-c**. **e-f**: Optimal reads per cell and number of total cells across TPM thresholds and FCs calculated from the two reference datasets. Similar to the setup in **Fig. 4d-e**. **g-h**: Optimal cost for different pilot dataset against different TPM analysis thresholds and effect sizes of interest while varying reads per cell and cells per target calculated from the two reference datasets. Similar to the setup in **Fig. 4f-g**.

### Extended Data Tables

**Extended Data Table 1.** PerturbPlan supports 11 design problems across 6 variables. For each of the six variables potentially involved in the optimization for each design problem, it is indicated what role the variable plays in the optimization (if any). Variables not indicated in the table (e.g., number of targets or multiplicity of infection) are kept fixed throughout.

|  | Power | Cost | Cells / target | Reads / cell | Expr. thresh. | Fold change | Example design problem |
| --- | --- | --- | --- | --- | --- | --- | --- |
| 1 | ≥ | N/A | min | = | = | = | Find the smallest cells/target to achieve at least 80% power to detect 15% FC for genes with at least 1 TPM, assuming 10k reads per cell. |
| 2 | ≥ | N/A | = | min | = | = | Find the smallest reads/target to achieve at least 80% power to detect 15% FC for genes with at least 1 TPM, assuming 500 cells/target. |
| 3 | ≥ | N/A | = | = | = | min | Find the smallest FC against which the power is at least 80% for genes with at least 1 TPM, assuming 500 cells/target and 10k reads/cell. |
| 4 | ≥ | N/A | = | = | min | = | Find the smallest expression threshold for which power to detect 15% FC is at least 80%, assuming 500 cells/target and 10k reads/cell. |
| 5 | ≥ | min | var | var | = | = | What is the minimum-cost combination of cells/target and reads/cell achieving at least 80% power to detect 15% FC for genes with at least 1 TPM? |
| 6 | ≥ | ≤ | var | var | = | min | Within a \$20k budget, what combination of cells/target and reads/cell give the minimum FC against which power is at least 80% for genes with at least 1 TPM? |
| 7 | ≥ | ≤ | var | = | = | min | Within a \$20k budget, what value of cells/target gives the minimum FC against which power is at least 80% for genes with at least 1 TPM, assuming 10k reads/cell? |
| 8 | ≥ | ≤ | = | var | = | min | Within a \$20k budget, what value of reads/cell gives the minimum FC against which power is at least 80% for genes with at least 1 TPM, assuming 500 cells/target? |
| 9 | ≥ | ≤ | var | var | min | = | Within a \$20k budget, what combination of cells/target and reads/cell give the smallest TPM threshold for which power to detect a 15% FC is at least 80%? |
| 10 | ≥ | ≤ | var | = | min | = | Within a \$20k budget, what value of cells/target gives the smallest TPM threshold for which power to detect a 15% FC is at least 80%, assuming 10k reads/cell? |
| 11 | ≥ | ≤ | = | var | min | = | Within a \$20k budget, what value of reads/cell gives the smallest TPM threshold for which power to detect a 15% FC is at least 80%, assuming 500 cells/target? |

