## Supplementary Note for "PerturbPlan: An analytical framework for designing Perturb-seq experiments"

### Supplementary Note accompanying "PerturbPlan: An analytical framework for designing Perturb-seq experiments"

May 15, 2026

### 1 Setup for power formula

#### 1.1 Notation

This notation mostly follows that of the Methods section, except that in the Methods section we do not use the superscripts "ref" for reference data to enhance readability.

**Experimental choices.** Let $L$ be the number of elements targeted, $K_0$ the number of non-targeting gRNAs, $K$ the number of gRNAs targeting each element, $N$ the total number of cells, $J$ the total number of genes captured, MOI the multiplicity of infection.

**Analysis choices.** Let $\mathcal{P} \subseteq \{1, \ldots, L\} \times \{1, \ldots, J\}$ be the element-gene pairs to test (after any expression filtering), and $q$ the FDR target level.

**Effect sizes.** Let $(\mu, \sigma^2)$ be the mean and variance of the fold changes (across gRNAs) in the expression of any gene upon perturbation of an element $l$ that regulates it. Let $\beta_{kl,j}$ denote the fold change in the expression of gene $j$ after perturbation by the $k$th gRNA targeting element $l$. Let $I_{lj} \in \{0, 1\}$ indicate whether element $l$ regulates gene $j$, and let $\pi$ be the proportion of pairs $\mathcal{P}$ where the element regulates the gene.

**Gene expression parameters.** Let $\alpha_j$ and $\theta_j$ denote the baseline relative expression (i.e., expected TPM / $10^6$) and dispersion, respectively, of the $j$th gene.

**Planned single-cell CRISPR screen data.** Let $R$ be the average number of mapped reads per cell, $S$ the average number of distinct *sequenced* UMIs per cell (library size), and $T$ the average number of distinct *captured* UMIs per cell, so that $S \leq T$ because not all captured UMIs are sequenced. Let $Y^t_{ikl,j}$ denote the UMI count of gene $j$ in the $i$-th cell receiving the $k$-th gRNA targeting element $l$ ("t" stands for "treatment"), and let $Y^c_{il,j}$ denote the expression count of gene $j$ in the $i$-th control cell for the comparison with element-gene pair $(l, j)$ ("c" stands for "control"). Let $N^t_{kl}$ be the number of cells receiving the $k$-th gRNA targeting element $l$ and $N^c_l$ the number of control cells for element $l$.

**Reference expression data.** Let $\{R^{\text{ref}}_u\}_u$ be the number of reads supporting each UMI $u$ in the reference expression data. Let $Y^{\text{ref}}_{ij}$ denote the UMI count of gene $j$ in the $i$-th cell, and let $N^{\text{ref}}$ be the total number of cells in the reference data.

#### 1.2 Data-generating model

We assume that whether element $l$ regulates gene $j$ is drawn from a Bernoulli prior distribution with success probability $\pi$, akin to the two-groups model (Efron, 2008):

$$I_{lj} \overset{\text{ind}}{\sim} \text{Ber}(\pi), \quad (l, j) \in \mathcal{P}. \tag{1}$$

Given these indicators, we model the effect sizes as follows:

$$\beta_{kl,j} = \begin{cases} 1, & I_{lj} = 0; \\ \text{drawn from } N(\mu, \sigma^2), & I_{lj} = 1, \end{cases} \quad k = 1, \ldots, K; \; (l, j) \in \mathcal{P}. \tag{2}$$

Given an average library size $S$ and the effect sizes $\beta_{kl,j}$, we model the per-cell gene expression counts using a negative binomial distribution:

$$Y^t_{ikl,j} \mid S, \beta_{kl,j}, \alpha_j, \theta_j \sim \text{NB}(S\alpha_j\beta_{kl,j}, \theta_j), \quad i = 1, \ldots, N^t_{kl}, \quad k = 1, \ldots, K; \tag{3}$$

$$Y^c_{il,j} \mid S, \alpha_j, \theta_j \sim \text{NB}(S\alpha_j, \theta_j), \quad i = 1, \ldots, N^c_l. \tag{4}$$

#### 1.3 Testing procedure under analysis

We analyze the following two-step procedure:

1. Apply the simplified `sceptre` test to get a $p$-value $p_{lj}$ for each pair $(l, j) \in \mathcal{P}$.
2. Apply the BH correction to the $p$-values obtained in step 1.

In the rest of this section, we walk through each of these two steps.

**1. Simplified `sceptre` test.** The simplified `sceptre` test statistic can be written

$$T_{lj} = \frac{\bar{Y}^t_{lj} - \bar{Y}^c_j}{\sqrt{\bar{Y}_j(1 + \bar{Y}_j/\theta_j)(1/N^t_l + 1/N^c_l)}}, \tag{5}$$

where

$$\bar{Y}^t_{lj} \equiv \frac{1}{N^t_l}\sum_{k=1}^{K}\sum_{i=1}^{N^t_{kl}} Y^t_{ikl,j}, \; \bar{Y}^c_j \equiv \frac{1}{N^c_l}\sum_{i=1}^{N^c_l} Y^c_{i,j}, \; \bar{Y}_j = \frac{N^t}{N^t + N^c}\bar{Y}^t_{lj} + \frac{N^c}{N^t + N^c}\bar{Y}^c_j. \tag{6}$$

Here, we define $N^t_l = \sum_{k=1}^{K} N^t_{kl}$. The $p$-value $p_{lj}$ based on the score test statistic (5) is computed by extracting a normal tail probability based on $T_{lj}$ and the sidedness of the test:

$$p_{lj} = \begin{cases} \Phi(T_{lj}), & \text{left-sided test;} \\ 1 - \Phi(T_{lj}), & \text{right-sided test;} \\ 2(1 - \Phi(|T_{lj}|)), & \text{two-sided test.} \end{cases} \tag{7}$$

**2. BH correction.** The BH correction applied to the $p$-values $\{p_{lj} : (l, j) \in \mathcal{P}\}$ leads to a final set of significant pairs

$$\mathcal{S} = \{(l, j) \in \mathcal{P} : p_{lj} \leq \hat{t}\},$$

where (Storey, Taylor, and Siegmund, 2004)

$$\hat{t} \equiv \max \left\{ t \in [0, 1] : \widehat{\text{FDP}}(t) \equiv \frac{|\mathcal{P}| \cdot t}{\sum_{(l,j)\in\mathcal{P}} \mathbb{1}(p_{lj} \leq t)} \leq q \right\}. \tag{8}$$

##### 1.4 Definition of power

Letting $\mathcal{P}_1 = \{(l, j) \in \mathcal{P} : I_{lj} = 1\}$ be the set of pairs with regulatory relationships, we define the power as the expected true positive proportion:

$$\text{Power} = \mathbb{E}\left[\frac{|\mathcal{S} \cap \mathcal{P}_1|}{|\mathcal{P}_1|}\right]. \tag{9}$$

Our goal is to approximate this notion of power for the procedure described in Section 1.3 under the data-generating model described in Section 1.2. We do so in the next section.

### 2 Derivation of power formula

#### 2.1 Step 1: Baseline expression summarization

In order to approximate the parameters $(\alpha_j, \theta_j)$ governing the expressions of each gene $j$, we fit an NB regression model to the reference expression data for each gene $j$:

$$Y_{ij}^{\text{ref}} \sim \text{NB}(S_i^{\text{ref}}\alpha_j, \theta_j), \quad i = 1, \ldots, N^{\text{ref}}. \tag{10}$$

Here, $S_i^{\text{ref}} = \sum_{j=1}^{J} Y_{ij}^{\text{ref}}$ is the library size of cell $i$ in the reference data. We fit $\alpha_j$ using the closed form MLE $\hat{\alpha}_j = \sum_{i=1}^{N^{\text{ref}}} Y_{ij}^{\text{ref}} / \sum_{i=1}^{N^{\text{ref}}} S_i^{\text{ref}}$. $\theta_j$ is also fit via the MLE, with initialization and fallback strategies as in `sceptre` (Barry et al., 2021).

#### 2.2 Step 2: Library size computation

We estimate the relationship between $S$, the average library size (number of distinct UMIs captured and sequenced per cell), and $R$, the average number of mapped reads per cell, from the empirical reads-per-UMI distribution $\{R_u^{\text{ref}}\}_u$ in a reference dataset using the `preseqR` package (Deng, Daley, and Smith, 2015). The `preseqR.rSAC()` function in this package, based on a combination of the zero-truncated negative binomial model and the rational function approximation approach (Daley and Smith, 2013), gives a curve $\hat{F}_{\text{tot}} : R_{\text{tot}} \mapsto S_{\text{tot}}$ mapping $R_{\text{tot}}$ (the number of total mapped reads) to $S_{\text{tot}}$ (the number of total UMIs represented among these reads). This curve is applicable to the library based on the number of cells, $N^{\text{ref}}$, in the reference expression data. However, we reasoned that this curve could be transferred to the per-cell level and applied to planned experiments with potentially different numbers of cells. In particular, we reasoned that in the homogeneous populations of cells often used for single-cell CRISPR screens, reads often are distributed roughly equally across cells, and that the saturation curves are roughly equal across cells. Under these assumptions, the per-cell saturation curve $\hat{F}_{\text{cell}} : R \mapsto S$ mapping $R$ (the number of mapped reads per cell) to $S$ (the number of UMIs per cell), inferred from the learned overall saturation curve $\hat{F}_{\text{tot}}$, is

$$\hat{F}_{\text{cell}}(R) = \frac{1}{N^{\text{ref}}} \hat{F}_{\text{tot}}(N^{\text{ref}} R), \tag{11}$$

where $R$ is the average number of mapped reads per cell. Given this per-cell saturation curve $\hat{F}_{\text{cell}}$, we obtain the average number of mapped reads per cell by multiplying the number of planned sequencing reads per cell by the mapping efficiency to get the average number of mapped reads per cell ($R$), and then computing $S = \hat{F}_{\text{cell}}(R)$ to get the average library size per cell.

#### 2.3 Step 3: Test statistic distributional approximation

The goal of this section is to obtain a normal approximation for the test statistic $T_{lj}$ (5), for each $(l, j) \in \mathcal{P}$:

$$T_{lj} \mid \{\beta_{kl,j}\}_k \dot{\sim} N(\eta_{lj}, \tau_{lj}^2), \tag{12}$$

where $\eta_{lj}$ and $\tau_{lj}^2$ denote the conditional mean and variance, respectively. They both depend on $\{\beta_{kl,j}\}_k, \alpha_j, \theta_j$.

**Approximations of cell counts.** We make the approximation that each gRNA is assigned to the same number of cells. Under these assumptions, we have for each $(k, l)$ that

$$N_{kl}^t = \frac{N \times \text{MOI}}{KL + K_0}$$

and

$$N_l^c = N^c = \begin{cases} N - N^t & \text{if using complement control cells;} \\ \frac{K_0}{KL+K_0} \times N \times \text{MOI} & \text{if using non-targeting control cells.} \end{cases}$$

Here, complement control cells for a given gRNA (used in high-MOI settings) are the set of cells not receiving the gRNA, whereas non-targeting control cells (often used in low-MOI settings) are the set of cells receiving non-targeting gRNAs.

**Approximation of the conditional mean.** By applying the law of large numbers (LLN) to numerator and denominator of $T_{lj}$ conditionally on $\{\beta_{kl,j}\}_k$, we have

$$\mathbb{E}[T_{lj}] \approx \frac{\mathbb{E}[\bar{Y}_{lj}^t] - \mathbb{E}[\bar{Y}_{lj}^c]}{\sqrt{\mathbb{E}[\bar{Y}_{lj}] \left(1 + \frac{\mathbb{E}[\bar{Y}_{lj}]}{\theta_j}\right) \left(\frac{1}{N^t} + \frac{1}{N^c}\right)}} \equiv \eta_{lj}. \tag{13}$$

Here and throughout this section, all expectations and variances are implicitly conditional on $\{\beta_{kl,j}\}_k$. From the data-generating model (3)–(4), we have:

$$\mathbb{E}[\bar{Y}_{lj}^t] = S\alpha_j\bar{\beta}_{lj},\ \mathbb{E}[\bar{Y}_{lj}^c] = S\alpha_j,\ \mathbb{E}[\bar{Y}_{lj}] = \frac{N^t}{N^t+N^c}S\alpha_j\bar{\beta}_{lj} + \frac{N^c}{N^t+N^c}S\alpha_j,$$

where $\bar{\beta}_{lj} = \frac{1}{K}\sum_{k=1}^K \beta_{kl,j}$ is the average of fold changes across the $K$ gRNAs targeting element $l$. This leads us to the formula

$$\eta_{lj} = \frac{S\alpha_j(\bar{\beta}_{lj}-1)}{\sqrt{\left[S\alpha_j\frac{N^t\bar{\beta}_{lj}+N^c}{N^t+N^c}\right]\left(1+\frac{S\alpha_j}{\theta_j}\frac{N^t\bar{\beta}_{lj}+N^c}{N^t+N^c}\right)\left(\frac{1}{N^t}+\frac{1}{N^c}\right)}} \tag{14}$$

**Approximation of the conditional variance.** By treating the denominator of $T_{lj}$ as nearly constant, since its fluctuation tends to be small compared to the numerator, we obtain

$$\mathrm{Var}[T_{lj}] \approx \frac{\mathrm{Var}[\bar{Y}_{lj}^t]+\mathrm{Var}[\bar{Y}_{lj}^c]}{\bar{Y}_{lj}\left(1+\frac{\bar{Y}_{lj}}{\theta_j}\right)\left(\frac{1}{N^t}+\frac{1}{N^c}\right)} \approx \frac{\mathrm{Var}[\bar{Y}_{lj}^t]+\mathrm{Var}[\bar{Y}_{lj}^c]}{\mathbb{E}[\bar{Y}_{lj}]\left(1+\frac{\mathbb{E}[\bar{Y}_{lj}]}{\theta_j}\right)\left(\frac{1}{N^t}+\frac{1}{N^c}\right)} \equiv \tau_{lj}^2, \tag{15}$$

where the second approximation $\bar{Y}_{lj} \approx \mathbb{E}[\bar{Y}_{lj}]$ is by the LLN. To compute $\tau_{lj}^2$, it remains to compute the variances of $\mathrm{Var}[\bar{Y}_{lj}^t]$ and $\mathrm{Var}[\bar{Y}_{lj}^c]$. To compute the variance of $\bar{Y}_{lj}^c$, recall from the data-generating model (4) that each control-cell expression is $\mathrm{NB}(S\alpha_j, \theta_j)$. Hence,

$$\mathrm{Var}[\bar{Y}_{lj}^c] = \frac{1}{N^c}\mathrm{Var}[Y_{il,j}^c] = \frac{1}{N^c}S\alpha_j\left(1+\frac{S\alpha_j}{\theta_j}\right).$$

Similarly, we can compute the variance for treated cells:

$$\begin{aligned}\mathrm{Var}[\bar{Y}_{lj}^t] &= \frac{1}{(N^t)^2}\sum_{k=1}^K\sum_{i=1}^{N_{kl}^t}\mathrm{Var}[Y_{ikl,j}^t] \\ &= \frac{1}{(N^t)^2}\sum_{k=1}^K\sum_{i=1}^{N_{kl}^t}S\alpha_j\beta_{kl,j}\left(1+\frac{S\alpha_j\beta_{kl,j}}{\theta_j}\right) \\ &= \frac{1}{N^t}S\alpha_j\bar{\beta}_{lj} + \frac{1}{N^t}\frac{S^2\alpha_j^2}{\theta_j}\cdot\frac{1}{K}\sum_{k=1}^K\beta_{kl,j}^2,\end{aligned}$$

where $\bar{\beta}_{lj} = \frac{1}{K}\sum_{k=1}^K \beta_{kl,j}$. The last equality is true because of the assumption $N^t = KN_{kl}^t$. Putting together the preceding three displays yields the final formula

$$\tau_{lj}^2 = \frac{\frac{S\alpha_j\bar{\beta}_{lj}}{N^t} + \frac{S^2\alpha_j^2}{N^t\theta_j}\left(\frac{1}{K}\sum_{k=1}^K\beta_{kl,j}^2\right) + \frac{S\alpha_j}{N^c}\left(1+\frac{S\alpha_j}{\theta_j}\right)}{\left[S\alpha_j\frac{N^t\bar{\beta}_{lj}+N^c}{N^t+N^c}\right]\left(1+\frac{S\alpha_j}{\theta_j}\frac{N^t\bar{\beta}_{lj}+N^c}{N^t+N^c}\right)\left(\frac{1}{N^t}+\frac{1}{N^c}\right)}. \tag{16}$$

#### 2.4 Step 4: BH cutoff approximation

Given the BH $p$-value threshold definition (8), we seek to approximate the function $\widehat{\mathrm{FDP}}(t)$ for each $t \in [0,1]$. To this end, note that in the case of left-sided tests (the other two cases are analogous), we have by the LLN that

$$\widehat{\mathrm{FDP}}(t) = \frac{t}{\frac{1}{|\mathcal{P}|}\sum_{(l,j)\in\mathcal{P}} \mathbb{1}(p_{lj} \leq t)} \approx \frac{t}{\frac{1}{|\mathcal{P}|}\sum_{(l,j)\in\mathcal{P}} \mathbb{P}[p_{lj} \leq t]}. \tag{17}$$

Note that, under the modeling assumptions 1.2 and the approximation that equal numbers of cells receive each gRNA, the distributions of the $p$-values $p_{lj}$ across elements $l$ are the same for a given gene $j$. Letting $P_j = \sum_{(l,j')\in\mathcal{P}} I(j' = j)$ be the number of pairs involving gene $j$ and $P = \sum_{j=1}^{J} P_j = |\mathcal{P}|$, we can therefore rewrite the preceding expression as

$$\widehat{\mathrm{FDP}}(t) \approx \frac{t}{\frac{1}{P}\sum_{j=1}^{J} P_j \mathbb{P}[p_{lj} \leq t]}, \tag{18}$$

where $l$ is the index of a generic element paired with gene $j$. We further derive that

$$\begin{aligned}
\widehat{\mathrm{FDP}}(t) &\approx \frac{t}{\frac{1}{P}\sum_{j=1}^{J} P_j \mathbb{P}[p_{lj} \leq t]} \\
&= \frac{t}{(1-\pi)t + \pi\frac{1}{P}\sum_{j=1}^{J} P_j \mathbb{P}[p_{lj} \leq t \mid I_{lj} = 1]} && \text{by model (1)} \\
&= \frac{t}{(1-\pi)t + \pi\frac{1}{P}\sum_{j=1}^{J} P_j \mathbb{E}[\mathbb{P}[\Phi(T_{lj}) \leq t \mid \{\beta_{kl,j}\}_k]|I_{lj} = 1]} && \text{by def. (7)} \\
&\approx \frac{t}{(1-\pi)t + \pi\frac{1}{P}\sum_{j=1}^{J} P_j \mathbb{E}[\Phi(\tau_{lj}^{-1}(\Phi^{-1}(t) - \eta_{lj}))|I_{lj} = 1]} && \text{by result (12)} \\
&\approx \frac{t}{(1-\pi)t + \pi\frac{1}{P}\sum_{j=1}^{J} P_j \Phi(\tau_{lj}^{-1}(\Phi^{-1}(t) - \eta_{lj}))} && \text{by LLN} \\
&\equiv \widetilde{\mathrm{FDP}}(t).
\end{aligned}$$

The quantity in the penultimate line is obtained by drawing one realization of $\beta_{kl,j} \overset{\text{i.i.d.}}{\sim} N(\mu, \sigma^2)$ for each $(k,j)$, as dictated by the model (2), and plugging it into the formulas for $\eta_{lj}$ (14) and $\tau_{lj}$ (16). To reduce computational cost in the case $J > 1000$, we further approximate the sum over $j$ by subsampling 1000 genes $\mathcal{J}$ from $\{1,\ldots,J\}$ without replacement and with weights $P_j/P$, and approximating

$$\frac{1}{P}\sum_{j=1}^{J} P_j \Phi(\tau_{lj}^{-1}(\Phi^{-1}(t) - \eta_{lj})) \approx \frac{1}{|\mathcal{J}|}\sum_{j\in\mathcal{J}} \Phi(\tau_{lj}^{-1}(\Phi^{-1}(t) - \eta_{lj})). \tag{19}$$

The approximation $\widetilde{\mathrm{FDP}}(t)$ for each $t$ gives rise to the following approximation to the BH cutoff:

$$\tilde{t} = \max\left\{t \in [0,1] : \widetilde{\mathrm{FDP}}(t) \leq q\right\}. \tag{20}$$

In practice, $\widetilde{\mathrm{FDP}}(t)$ is an increasing function of $t$ (we provide theoretical support for this observation in Appendix A), so we find $\tilde{t}$ by solving the equation $\widetilde{\mathrm{FDP}}(t) = q$ via bisection.

#### 2.5 Step 5: Overall power calculation

Recalling the definition of power (9) and again taking the case of left-sided tests as a representative example, we find that

$$\begin{aligned}
\text{Power} &= \mathbb{E}\left[\frac{\mathcal{S}\cap\mathcal{P}_1}{|\mathcal{P}_1|}\right] \\
&\approx \frac{1}{\pi|\mathcal{P}|}\mathbb{E}[|\mathcal{S}\cap\mathcal{P}_1|] \\
&= \frac{1}{\pi|\mathcal{P}|}\sum_{j=1}^{J} P_j \mathbb{P}[p_{lj}\leq \hat{t}, I_{lj}=1] \\
&= \frac{1}{|\mathcal{P}|}\sum_{j=1}^{J} P_j \mathbb{P}[p_{lj}\leq \hat{t}\mid I_{lj}=1] \\
&\approx \frac{1}{|\mathcal{P}|}\sum_{j=1}^{J} P_j \Phi\left(\tau_{lj}^{-1}(\Phi^{-1}(\tilde{t})-\eta_{lj})\right).
\end{aligned} \tag{21}$$

In the first step, we used the fact that $|\mathcal{P}_1|\approx\pi|\mathcal{P}|$ by the LLN. In the last step, we used the approximations $\hat{t}\approx\tilde{t}$ and

$$\frac{1}{|\mathcal{P}|}\sum_{j=1}^{J} P_j \mathbb{P}[p_{lj}\leq t\mid I_{lj}=1]\approx\frac{1}{|\mathcal{P}|}\sum_{j=1}^{J} P_j \Phi\left(\tau_{lj}^{-1}(\Phi^{-1}(t)-\eta_{lj})\right),$$

both derived in Step 4. If $J>1000$, we replace the sum in the last step of the formula (21) by its subsampled analog based on the genes selected in Step 4.

### 3 Comparison of PerturbPlan to scPower

scPower (Schmid et al., 2021) is an experimental design and power analysis tool for single-cell transcriptomic studies, providing guidance for differential gene expression and expression quantitative trait loci analyses. Because PerturbPlan is also designed to support experimental design in a different differential expression context, we conduct a systematic comparison between PerturbPlan and scPower. We first describe the features shared by the two methods (Section 3.1), and then highlight six areas in which PerturbPlan departs from scPower (Section 3.2).

#### 3.1 Overlapping features of scPower and PerturbPlan

Both tools provide experimental design guidance for differential expression analyses and develop analytical power formulas under a negative binomial model. Moreover, both tools adopt an analytical approximation to the BH cutoff based on the empirical process perspective of the BH procedure (Genovese and Wasserman, 2002). They also incorporate sequencing saturation modeling to capture the relationship between sequencing depth and library size. In addition, both scPower and PerturbPlan account for the trade-off between sequencing cost and library preparation cost, and offer interactive web applications.

#### 3.2 Distinct features of PerturbPlan

We now discuss the distinct features of PerturbPlan compared to scPower.

1. **CRISPR-specific experimental parameters in web application:** PerturbPlan's web application is specifically tailored for single-cell CRISPR screen experimental design, including a number of variables not applicable to scPower's setting and therefore not present in their web application: multiplicity of infection (MOI), number of perturbation targets, number of gRNAs per target, number of non-targeting gRNAs, and gRNA-level variability, among others.

2. **CRISPR-specific variables in analytical power formula:** PerturbPlan's power formula explicitly incorporates CRISPR-specific variables. While some variables could potentially be mapped to scPower's framework, others are unique to CRISPR screens. For example, PerturbPlan models the number of gRNAs per element and the variability in gRNA perturbation efficiency, which have no equivalent in scPower's power formula.

3. **More accurate saturation curve fitting:** PerturbPlan employs a preseqR-based sequencing saturation curve fit that models the molecular sampling process inherent in sequencing (see Section 2.2). By contrast, scPower employs an ad hoc log-linear regression of UMI counts on log-transformed read counts. When applied to real Perturb-seq datasets, the preseqR-based saturation curve provides improved goodness of fit relative to the scPower fit (Extended Data Fig. 3a-b).

4. **Wide span of supporting experimental design problems:** PerturbPlan supports 11 distinct experimental design problems commonly encountered in single-cell CRISPR screens (Extended Data Table 1). These include complex optimization tasks, such as the following: Find the smallest fold change that can be detected among perturbation-gene pairs involving genes with at least 5 TPM, with at least 80% power, for under $10,000, allowing cells/target and reads/cell to vary. By contrast, scPower supports a more limited set of experimental design scenarios, primarily focusing on optimizing the number of cells and/or sequencing depth.

5. **Flexible and interactive slider functionality:** The PerturbPlan app provides an interactive interface that allows users to explore a range of design settings through sliders controlling up to six key parameters and a visualization that includes all parameter settings pinned by the user (Figure 1g, Extended Data Figure 2). This functionality enables users to see how varying parameters impacts optimal experimental designs and their associated costs. By contrast, the scPower application does not provide such functionality.

6. **Ready-to-use reference expression data for common cell types:** While both scPower and PerturbPlan support preprocessing of arbitrary reference expression datasets, PerturbPlan further lowers the barrier to experimental planning by providing curated, ready-to-use reference expression data for cell types that are most commonly used in CRISPR-based studies: K562, A549, iPSCs, iPSC-derived neurons, CD8+ T cells, and THP-1 cells.

### 4 Simulation benchmarking details

To validate the analytical power formula presented in the main text, we conducted a simulation-based benchmarking study using the `sceptre` package (Barry et al., 2021). The simulation pipeline is publicly available at https://github.com/ZiangNiu6/Sceptre_Power_Simulations. This repository is a frozen fork of the simulation pipeline originally developed by James Galante (https://github.com/jamesgalante/Sceptre_Power_Simulations); the core data-generation and testing code is unchanged, while the fork pins the exact parameter sweeps, cluster configuration, and aggregation scripts used to produce the results reported here, ensuring full reproducibility independent of upstream development. Below we describe the simulation setup, data generation procedure, and power computation methodology.

#### 4.1 Simulation overview

The simulation adopts a semi-synthetic approach: it draws on real single-cell CRISPR screen data to calibrate distributional parameters (gene-level dispersions, cell-level size factors, and baseline mean expression levels) while generating synthetic perturbation assignments and count matrices under controlled effect sizes. Power is then estimated as the fraction of simulation replicates in which a perturbation effect is detected at a given significance level.

#### 4.2 Parameter sweep

The simulation pipeline is organized as two separate one-dimensional parameter sweeps:

- **Effect size sweep**: 8 values of $\delta$ ranging from 0.05 to 0.40 in increments of 0.05, representing the fractional decrease in gene expression due to a perturbation (fold-change $= 1 - \delta$), with the number of treated cells per perturbation fixed at 1,000.
- **Sample size sweep**: 8 values of the number of treated cells per perturbation ranging from 250 to 2,000 in increments of 250, with the effect size fixed at $\delta = 0.15$.

Each parameter setting is run as an independent Snakemake workflow on a computing cluster.

#### 4.3 Data generation

For each simulation replicate, the data generation proceeds as follows:

**Baseline calibration.** A pre-existing single-cell experiment (SCE) object provides the real-data backbone. Gene-level dispersions and mean expression levels are estimated from the real count matrix using DESeq2 with the "poscounts" size factor method and parametric dispersion estimation. Cell-level size factors are resampled with replacement from the empirical distribution.

**Guide assignment.** For each perturbation target, the number of cells assigned to each guide RNA is drawn from a Poisson distribution with mean $N_t/K$, where $N_t$ is the number of treated cells for that target and $K$ is the number of guides per target. Cells are randomly selected without replacement from the pool. Each cell is assigned to exactly one guide.

**Effect size matrix construction.** For each gene–perturbation pair, guide-level fold-changes for targeting guides are drawn from $\mathcal{N}(1-\delta,\, \sigma^2_{\text{guide}})$ with $\sigma_{\text{guide}} = 0.13$, capturing guide-to-guide variability; fold-changes for non-targeting (control) guides are drawn from $\mathcal{N}(1,\, \sigma^2_{\text{guide}})$. The fold-changes are then centered so that the mean fold-change for perturbed cells equals $1-\delta$ exactly and the mean for control cells equals 1. Negative fold-changes are clipped to zero.

**Count simulation.** The simulated count for gene $j$ in cell $i$ is drawn from a negative binomial distribution:

$$Y_{ij} \sim \text{NB}\big(\mu_{ij},\, \phi_j^{-1}\big)\,, \tag{22}$$

where $\mu_{ij} = \bar{\mu}_j \cdot s_i \cdot e_{ij}$, $\bar{\mu}_j$ is the baseline mean expression for gene $j$, $s_i$ is the size factor for cell $i$, $e_{ij}$ is the effect size for gene $j$ in cell $i$ (equal to 1 for unperturbed cells), and $\phi_j$ is the DESeq2-estimated dispersion for gene $j$.

**Covariate resampling.** Cell-level covariates (number of nonzero features per cell and total UMI count per cell) are resampled with replacement from the real data to preserve realistic covariate distributions.

#### 4.4 Statistical testing

Each simulated dataset is analyzed using the `sceptre` discovery analysis pipeline with the following settings:

- **Side**: left-sided test (testing for decrease in expression).
- **Control group**: complement (all non-perturbed cells serve as controls).
- **gRNA integration strategy**: union (a cell is considered perturbed if it contains any targeting guide).
- **Resampling mechanism**: permutations with skew-normal approximation.
- **Multiplicity of infection**: high MOI.
- **gRNA assignment**: thresholding with threshold = 1.

#### 4.5 Power computation

For each gene–perturbation pair and each parameter configuration, 200 independent simulation replicates are run. Power is computed as the fraction of replicates in which the perturbation effect is detected after multiple testing correction. To account for the realistic setting in which only a small fraction of tested pairs are true positives, we adopt the following procedure for each replicate:

1. Let $n$ denote the number of true positive pairs tested in the simulation. Generate $n_{\text{null}} = n \cdot (1-\pi)/\pi$ null p-values drawn independently from Uniform$(0, 1)$, where $\pi = 0.05$ is the assumed proportion of true positives.

2. Combine the $n$ real p-values with the $n_{\text{null}}$ null p-values and apply the Benjamini–Hochberg procedure to the combined set.

3. Retain the adjusted p-values corresponding to the real pairs.

A pair is declared a discovery if its adjusted p-value falls below $\alpha = 0.1$. The power for each pair is then the fraction of the 200 replicates in which a discovery is made.

### A Monotonicity of $\widetilde{\mathrm{FDP}}(t)$

In this section, we prove that $\widetilde{\mathrm{FDP}}(t)$ is an increasing function of $t$ under the assumptions that $\tau_{lj}^2 = 1$ and that the sign of $\eta_{lj}$ aligns with the test direction. The former is a decent approximation in cases when the signal strength is not too large, and the latter is a basic power analysis assumption.

The FDP estimates for left-sided, right-sided, and two-sided tests can all be expressed as

$$\widetilde{\mathrm{FDP}} = \frac{t}{(1-\pi)t + \pi\frac{1}{|\mathcal{P}|}\sum_{j=1}^{J} P_j P_{lj}(t)} = \frac{1}{(1-\pi)1 + \pi\frac{1}{|\mathcal{P}|}\sum_{j=1}^{J} P_j \frac{P_{lj}(t)}{t}}, \tag{23}$$

where

$$P_{lj}(t) = \begin{cases} \Phi(\tau_{lj}^{-1}(\Phi^{-1}(t) - \eta_{lj})) & \text{left-sided test} \\ \Phi(\tau_{lj}^{-1}(\eta_{lj} - \Phi^{-1}(1-t))) & \text{right-sided test} \\ \Phi(\tau_{lj}^{-1}(\Phi^{-1}(t/2) - \eta_{lj})) + \Phi(\tau_{lj}^{-1}(\Phi^{-1}(t/2) + \eta_{lj})) & \text{two-sided test} \end{cases}. \tag{24}$$

Under the assumption that $\tau_{lj}^2 = 1$, it suffices to show that the following three functions are all decreasing in $t$:

$$g_1(t) = \frac{\Phi(\Phi^{-1}(t) - x)}{t}, \quad \text{for any } x \leq 0$$
$$g_2(t) = \frac{\Phi(x - \Phi^{-1}(1-t))}{t}, \quad \text{for any } x \geq 0$$
$$g_3(t) = \frac{\Phi(\Phi^{-1}(t/2) - x) + \Phi(\Phi^{-1}(t/2) + x)}{t}, \quad \text{for any } x \in \mathbb{R}.$$

In preparation to verify that these functions are decreasing, we first establish two properties of the *Mills ratio* $M(z) = \Phi(z)/\phi(z)$, which arises in the derivatives of $g_1(t), g_2(t), g_3(t)$.

#### A.1 Properties of the Mills ratio

We first compute the first and second derivatives of Mills ratio $M(z)$.

$$M'(z) = \frac{\phi^2(z) + z\Phi(z)\phi(z)}{\phi^2(z)} = \frac{\phi(z) + z\Phi(z)}{\phi(z)}. \tag{25}$$

The second derivative is given by

$$M''(z) = \frac{\Phi(z)\phi(z) + z\phi(z)(\phi(z) + z\Phi(z))}{\phi^2(z)} = \frac{\Phi(z) + z\phi(z) + z^2\Phi(z)}{\phi(z)}. \tag{26}$$

**The Mills ratio is increasing.** We now prove the derivative $M'(z)$ in (25) is positive. We compute the derivative of the numerator in (25):

$$-z\phi(z) + z\phi(z) + \Phi(z) = \Phi(z) > 0.$$

Then when $z \to -\infty$, we know $\phi(z) + z\Phi(z)$ converges to 0. Therefore, we know $M'(z) > 0$ for any $z \in \mathbb{R}$. This shows that $M(z)$ is an increasing function.

**The Mills ratio is convex.** Now we show that the numerator in (26) is positive. We compute the derivative of the numerator:

$$\phi(z) + \phi(z) - z^2\phi(z) + z^2\phi(z) + 2z\Phi(z) = 2\phi(z) + 2z\Phi(z) \geq 0.$$

Since $\Phi(z) + z\phi(z) + z^2\Phi(z)$ converges to 0 as $z$ tends to $-\infty$, then we know $\Phi(z) + z\phi(z) + z^2\Phi(z)$ is positive for any $z \in \mathbb{R}$. This shows that $M''(z) > 0$ for any $z \in \mathbb{R}$, which implies that $M(z)$ is a convex function.

#### A.2 Proofs that $g_1, g_2, g_3$ are decreasing

$g_1(t)$**.** Write $t = \Phi(z)$ so that it is sufficient to show $f_x(z) = \Phi(z-x)/\Phi(z)$ is a decreasing function for any fixed $x \leq 0$. To see this, we compute the derivative

$$f_x'(z) = \frac{\phi(z-x)\phi(z)}{\Phi^2(z)}[M(z) - M(z-x)], \tag{27}$$

which is non-positive since $x \leq 0$ and $M(z)$ is an increasing function.

$g_2(t)$**.** Write $1 - t = \Phi(z)$ so that it is sufficient to show $f_x(z) = \Phi(x-z)/\Phi(-z)$ is an increasing function for any fixed $x \geq 0$. To see this, we compute

$$f_x'(z) = \frac{\phi(-z)\phi(x-z)}{\Phi^2(-z)}(M(x-z) - M(-z)),$$

which is non-negative since $x \geq 0$ and $M$ is an increasing function.

$g_3(t)$**.** We can write $t/2 = \Phi(z)$ so that it is sufficient to show $f_x(z) = \Phi(z-x)/\Phi(z) + \Phi(z+x)/\Phi(z)$ is a decreasing function on $z \in (-\infty, 0)$ for any fixed $x \in \mathbb{R}$. To see this, we compute

$$f_x'(z) = \frac{\Phi(z)(\phi(z-x) + \phi(z+x)) - \phi(z)(\Phi(z-x) + \Phi(z+x))}{\Phi^2(z)}.$$

If we define $\bar{M}_x(z) \equiv (M(z-x)\phi(z-x) + M(z+x)\phi(z+x))/(\phi(z+x) + \phi(z-x))$, then we can write

$$f_x'(z) = \frac{\phi(z)(\phi(z-x) + \phi(z+x))}{\Phi^2(z)} \cdot [M(z) - \bar{M}_x(z)].$$

We just need to prove that $M(z) - \bar{M}_x(z) \leq 0$ for any $x \in \mathbb{R}$. We take cases based on the sign of $x$. If $x = 0$, the conclusion is straightforward because $\bar{M}_0(z) = M(z)$. If $x > 0$, then since $M(z+x) \geq M(z) \geq M(z-x)$ due to the increasing property of the Mills ratio and $\phi(z+x) > \phi(z) > \phi(z-x)$, we find

$$\begin{aligned}\bar{M}_x(z) &= \frac{\phi(z-x)}{\phi(z-x) + \phi(z+x)}M(z-x) + \frac{\phi(z+x)}{\phi(z-x) + \phi(z+x)}M(z+x) \\ &\geq \frac{1}{2} \cdot (M(z-x) + M(z+x)) \geq M(z).\end{aligned}$$

The last inequality is due to Jensen's inequality, guaranteed by the convexity of Mills ratio. Finally, if $x < 0$, then $M(z + x) < M(z) < M(z - x)$ due to the increasing property of Mills ratio and $\phi(z + x) < \phi(z) < \phi(z - x)$ give

$$\begin{aligned}\bar{M}_x(z) &= \frac{\phi(z-x)}{\phi(z-x)+\phi(z+x)}M(z-x) + \frac{\phi(z+x)}{\phi(z-x)+\phi(z+x)}M(z+x) \\ &\geq \frac{1}{2}\cdot\left(M(z-x)+M(z+x)\right) \geq M(z).\end{aligned}$$

The last inequality is due to the convexity of Mills ratio.

#### A.3 Empirical evidence of monotonicity

To illustrate the practical relevance of our theoretical results, we numerically evaluate the function $\widetilde{\mathrm{FDP}}(t)$ using the real-data example shown in Figure 3 of the main text. Specifically, we consider two downsampling-based validation setups from the Gasperini dataset and plot $\widetilde{\mathrm{FDP}}(t)$ in Figure 1. We then compare the resulting FDP curves obtained using the minimum versus the median effect size estimates derived from the effect size distribution of the Gasperini data. As in the main text, we examine two effect size distributions with different ranges. Left-sided test is considered. In both panels of Figure 1, we observe that $\widetilde{\mathrm{FDP}}(t)$ increases monotonically as the significance cutoff $t$ becomes more liberal, consistent with our theoretical findings.

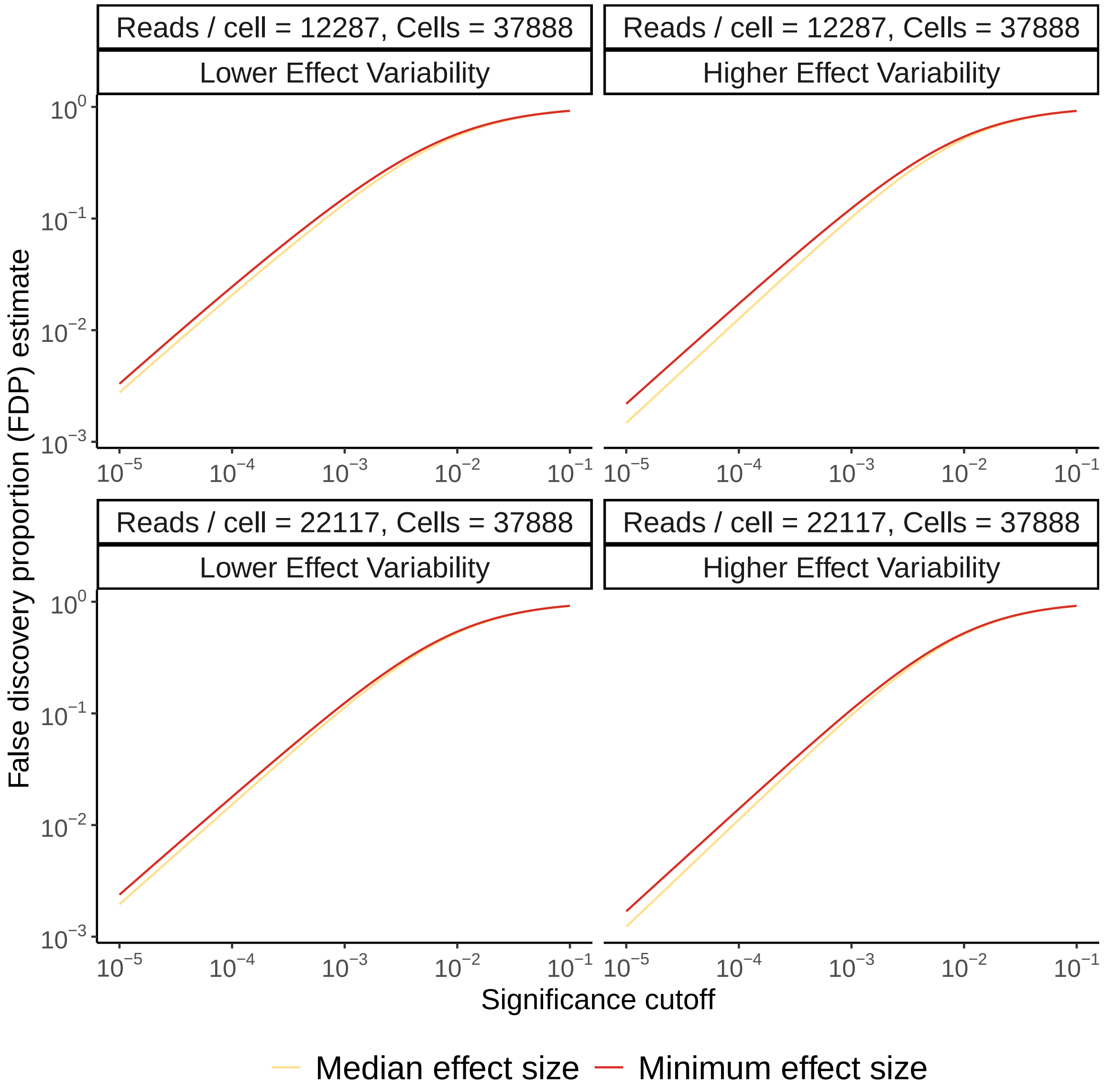


Figure 1: Empirical evaluation of the function $\widetilde{\mathrm{FDP}}(t)$ using two downsampling Gasperini data validation setups. Left-sided test is considered. The top panel corresponds to the setup with $37,888$ cells and $12,287$ reads per cell, while the bottom panel corresponds to the setup with $37,888$ cells and $22,117$ reads per cell. In both panels, we observe that the function $\widetilde{\mathrm{FDP}}(t)$ is monotonically increasing in significance cutoff $t$.
